# Signed Dopaminergic Asymmetry Tracks Contralateral Motor Laterality in Parkinson’s Disease

**DOI:** 10.64898/2026.09.23.753983

**Authors:** Harsh Milind Tirhekar, Priyanshi Yadav, Chandrajit Bajaj

**Affiliations:** Department of Computer Science, and Oden Institute for Computational Engineering and Sciences, The University of Texas at Austin, Austin, TX, USA; Department of Biomedical Engineering, National Institute of Technology Raipur, Raipur, Chhattisgarh, India; Department of Computer Science, Oden Institute for Computational Engineering and Sciences, The University of Texas at Austin, Austin, TX, USA

## Abstract

Parkinson’s disease is asymmetric, but representations that collapse left–right measurements discard that direction. We separated overall dopamine-transporter availability from hemispheric distribution: mean putamen specific-binding ratio (SBR) indexed overall DaT signal, whereas signed asymmetry retained which hemisphere was relatively more affected. Signed asymmetry aligned with contralateral MDS-UPDRS Part III laterality in PPMI (n=1,367; r=0.676) and S4 (n=56; r=0.551). In 1,204 PPMI landmarks, each 0.1 asymmetry corresponded to a 0.0507 increase in 12-month laterality (95% CI 0.0387–0.0626); held-out *R*^2^ improved 0.0176 over non-directional imaging but was uncertain under a bounded link. Exploratory imaging–clinical disagreement marked lower side persistence (OR=0.491 per IQR-standardized unit; 0.297–0.711). Signed direction therefore distinguishes mirror-image and conflicting dopaminergic–motor states erased by mean-only summaries. This is a reproducible biological coordinate, not global progression, treatment benefit or clinical utility.

## 1 Introduction

Clinical diagnosis of Parkinson’s disease (PD) is anchored in motor signs, but its biological organization is neither uniform nor symmetric [1]. Bradykinesia, rigidity and tremor often begin on one side of the body, and the initially more affected side can remain clinically dominant after symptoms become bilateral. The anatomical counterpart is a predominantly contralateral nigrostriatal dopaminergic deficit. A lower dopamine-transporter signal in the left putamen, for example, is expected to align with greater motor burden on the right side of the body [2, 3]. This directional relation is one of the most intuitive bridges between in-vivo neuroimaging and the neurological examination.

That bridge is easily weakened during analysis. Many studies replace left and right measurements with a mean, minimum, absolute difference or unsigned asymmetry index. Those summaries answer useful questions about overall level or degree of imbalance, but they cannot preserve which hemisphere is more affected. Two participants can have the same total MDS-UPDRS Part III score and the same absolute imaging asymmetry while carrying opposite anatomical and clinical directions. Once the sign is discarded, the model cannot distinguish those states.

For movement-disorder clinicians, neuroimaging scientists and PD-heterogeneity researchers, three quantities must therefore remain distinct. Lower mean bilateral putamen SBR indicates greater overall dopaminergic deficit; absolute asymmetry indicates the degree of left–right imbalance; and signed asymmetry identifies the hemisphere carrying the relatively greater deficit. The model hierarchy tests this third quantity against matched representations of overall DaT signal and unsigned imbalance. We use it as a spatial coordinate of nigrostriatal dysfunction, not as whole-disease severity, molecular etiology, alpha-synuclein stage or causal mechanism.

For a clinician or researcher reading a longitudinal record, this is not a minor encoding choice. The relevant question is not only whether dopaminergic signal and motor burden are abnormal, but whether the record still shows which hemisphere and body side are involved, when each source was measured, whether imaging and examination agree, and how much uncertainty remains about the future. A point prediction can estimate an outcome while hiding all four distinctions. A direction-aware state makes them visible.

The prognostic meaning of asymmetry has consequently remained unsettled. Cross-sectional studies have repeatedly linked lateralized striatal dopamine loss to contralateral motor expression [2, 3]. A PPMI study of 249 right-handed participants with greater than 20% putamen asymmetry reported different motor and cognitive trajectories according to the predominantly affected hemisphere [4]. In contrast, an analysis of 423 de-novo PPMI participants found that clinical and DaT-SPECT asymmetry did not predict one-, three- or five-year change in total MDS-UPDRS Parts I, II or III [5]. These findings are not necessarily contradictory. One asks whether hemispheric direction remains visible in future side-specific motor expression; the other asks whether the degree of asymmetry predicts how much the disease worsens globally.

This distinction matters as PD research moves toward biological definitions and multimodal prediction. NSD-ISS anchors neuronal alpha-synuclein disease in alpha-synuclein pathology and dopaminergic dysfunction, while SynNeurGe organizes disease through synuclein, neurodegeneration and genetic dimensions [6, 7]. Data-driven studies further combine clinical scales, DaT-SPECT, fluid biomarkers, genetics, cognition and digital measures to infer subtypes or forecast progression [8–13]. The scientific opportunity is substantial, but a larger feature matrix does not guarantee a more faithful patient state. Direction, measurement time, medication context, missingness and the unit of validation can matter more than model complexity [14].

Recent DaT-SPECT subtype-and-stage inference further demonstrates that regional dopaminergic patterns can organize prospective PD heterogeneity [13]. That work learns multiregional subtype and stage structure. Our question is deliberately different: we prespecify one signed anatomical coordinate, test whether its contralateral physiological meaning replicates without cohort-specific reorientation, and then ask whether it contributes future side-balance information beyond current clinical laterality and otherwise matched non-directional imaging.

Two earlier preprints from our group developed broad multiscale stratification and posterior-calibrated motor-state representations [15, 16]. The present study does not introduce another omnibus subtype or claim a calibrated mechanism probability. Instead, it isolates one falsifiable component of a mechanism-aware state: whether preserving the sign of dopaminergic anatomy adds future side-specific information after current clinical laterality, total motor burden, overall putamen binding and treatment context are represented.

We organized the test around claim-specific validity checks. First, we required the signed putamen-to-motor relation to reproduce in PPMI and the independent Systemic Synuclein Sampling Study (S4) [17]. Second, we prespecified a one-landmark-per-participant PPMI protocol before inspecting the longitudinal result. Third, we compared a direction-aware model with otherwise matched clinical, overall-DaT and magnitude-only models using participant-held-out, site-disjoint and strict-calendar evaluations. Fourth, we challenged the result with a non-lateralized axial outcome, an unnormalized right-minus-left outcome, exact scan-examination matching and an alternative cohort requiring identical explicit medication states. Finally, we tested whether imaging direction agreed with the historically affected side and whether onset side, handedness or incomplete paired motor items explained the association.

Our hypothesis was narrow. Signed putamen asymmetry would retain a positive association with future signed motor laterality and could improve continuous laterality prediction over the same imaging model without direction. Association and prediction were evaluated separately: a coherent coefficient need not yield useful incremental prediction. We did not hypothesize that direction would predict global disease progression, improve a binary affected-side decision, identify a causal mechanism or support treatment choice. Keeping those questions separate turns an apparently small encoding choice into a direct test of whether a multimodal representation preserves the biology it is intended to describe.

## 2 Results

### 2.1 Actual records make the directional state visible

Figure 1 begins with two actual patients whose lower-binding putamen lies in opposite hemispheres and whose greater motor burden lies on the corresponding contralateral body side. Their total MDS-UPDRS Part III scores are 31 and 12: they are opposite concordant configurations, not a severity-matched pair. The records retain source dates, treatment documentation and subsequent examinations, showing both what the integrated state resolves and where the forecast remains wrong. These examples were selected from a 32-record pool without their future outcomes or forecast errors; the selection and all four cases are described below.

**Figure 1:**
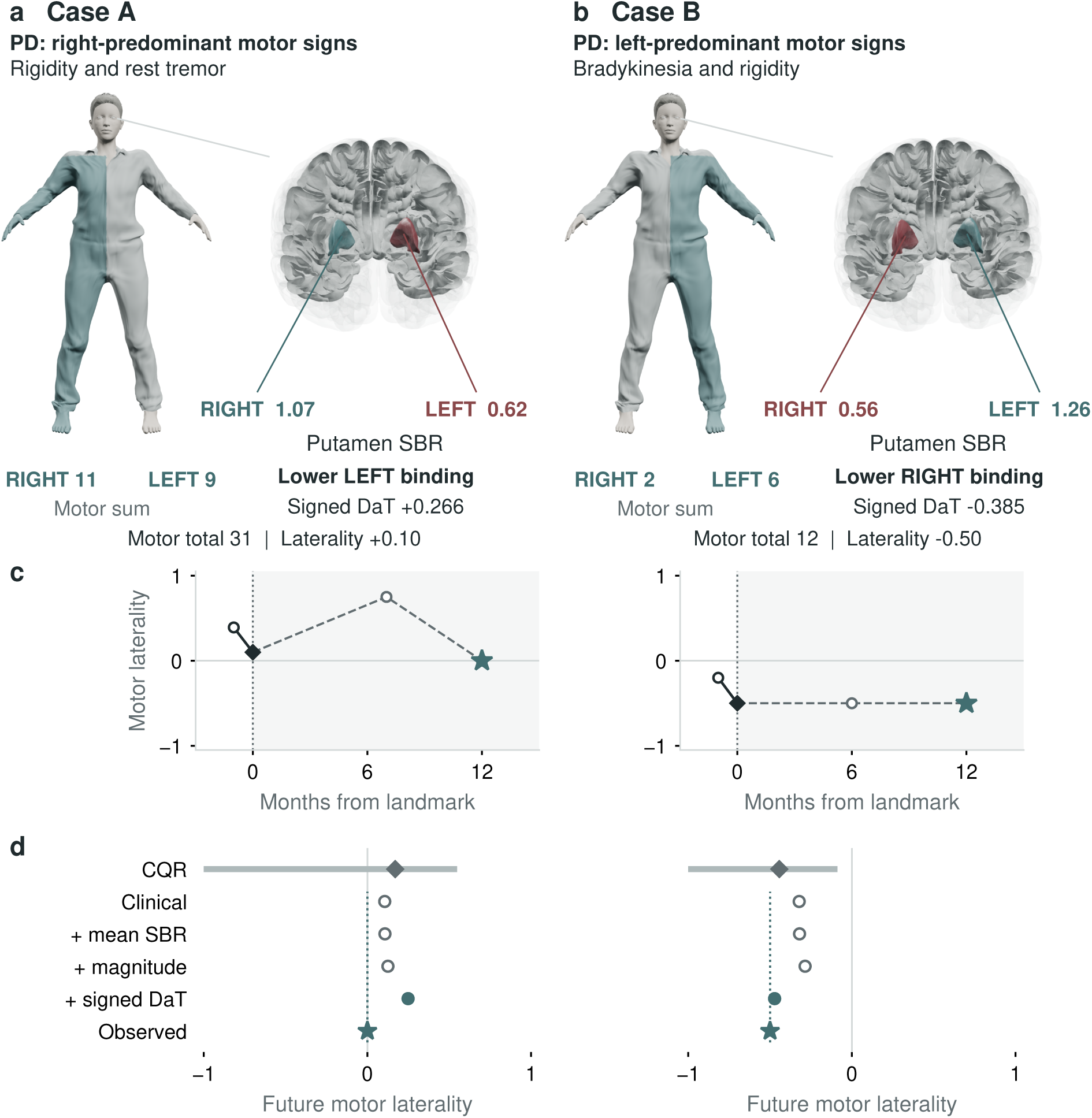
Observed motor phenotypes connect to opposite putamen states. a–b, LEFT/RIGHT labels identify regional SBR and paired motor scores. A has right-predominant rigidity/rest-tremor findings; B has left-predominant bradykinetic/rigid findings. These are item-based descriptions, not formal subtypes. Totals 31/12 are not severity-matched. Tint indicates aggregate body burden and relatively lower binding, not normative severity. Reference surfaces are not participant scans or motion. Medication state is unrecorded; LEDD is 100 mg/day. c, Examination summaries (circles), landmarks (diamonds), and selected future observations (stars); shading marks follow-up. d, Four held-out ridge forecasts; the star/dotted line marks observed laterality. The separate conformalized quantile regression (CQR) diamond and 90% interval belong only to the quantile model. Signed imaging worsens A’s error and improves B’s. Assets: MakeHuman/MPFB (CC0); NIH 3D Human Reference Atlas (CC BY 4.0).

The examined phenotype is more specific than the side sum alone. At Case A’s landmark, right/left arm-rigidity scores were 2/1 and arm rest-tremor scores were 1/0. In Case B, left/right finger-tapping and hand-movement scores were both 1/0. These are observed examination items, not symptoms inferred from an imaging score. All 11 paired domains, including findings that do not favor the dominant side, are retained in Supplementary Figure 7. The visual linkage therefore locates a measured motor pattern within a biologically compatible spatial state without claiming that regional binding explains every examination item.

### 2.2 Cohort roles keep biological replication separate from prognosis

Table 1 fixes the role of every cohort and sensitivity before the results are interpreted. Counts from distinct rows are overlapping analysis sets and must not be added into a single “multimodal sample size.”

**Table 1:**
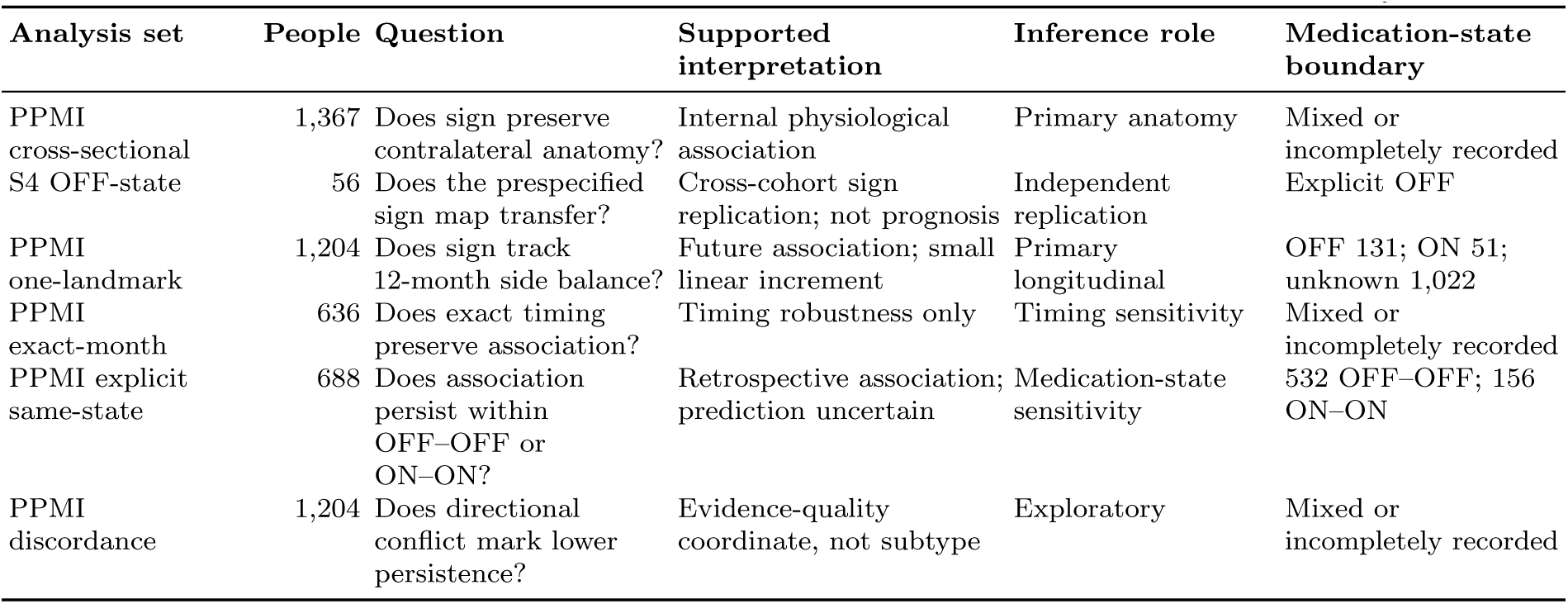
Cohort roles and inference boundaries in the direction-aware study.

### 2.3 A direction-aware record keeps anatomy, time and intended use visible

The computational chain is deliberately short. The measured left/right binding values in Figure 1a–b become a signed ratio; the 11 paired examination domains become a separate clinical ratio. A positive imaging ratio means lower left binding, whereas a positive clinical ratio means greater right-body burden. Their agreement is therefore contralateral anatomical consistency, not identity of the quantities. The landmark construction in Figure 2b places both observations before the outcome; matched ridge models then test whether keeping the imaging sign adds information beyond the clinical state and non-directional imaging. Panel d of Figure 1 shows the individual outputs, not guarantee that an anatomically coherent state will persist.

**Figure 2:**
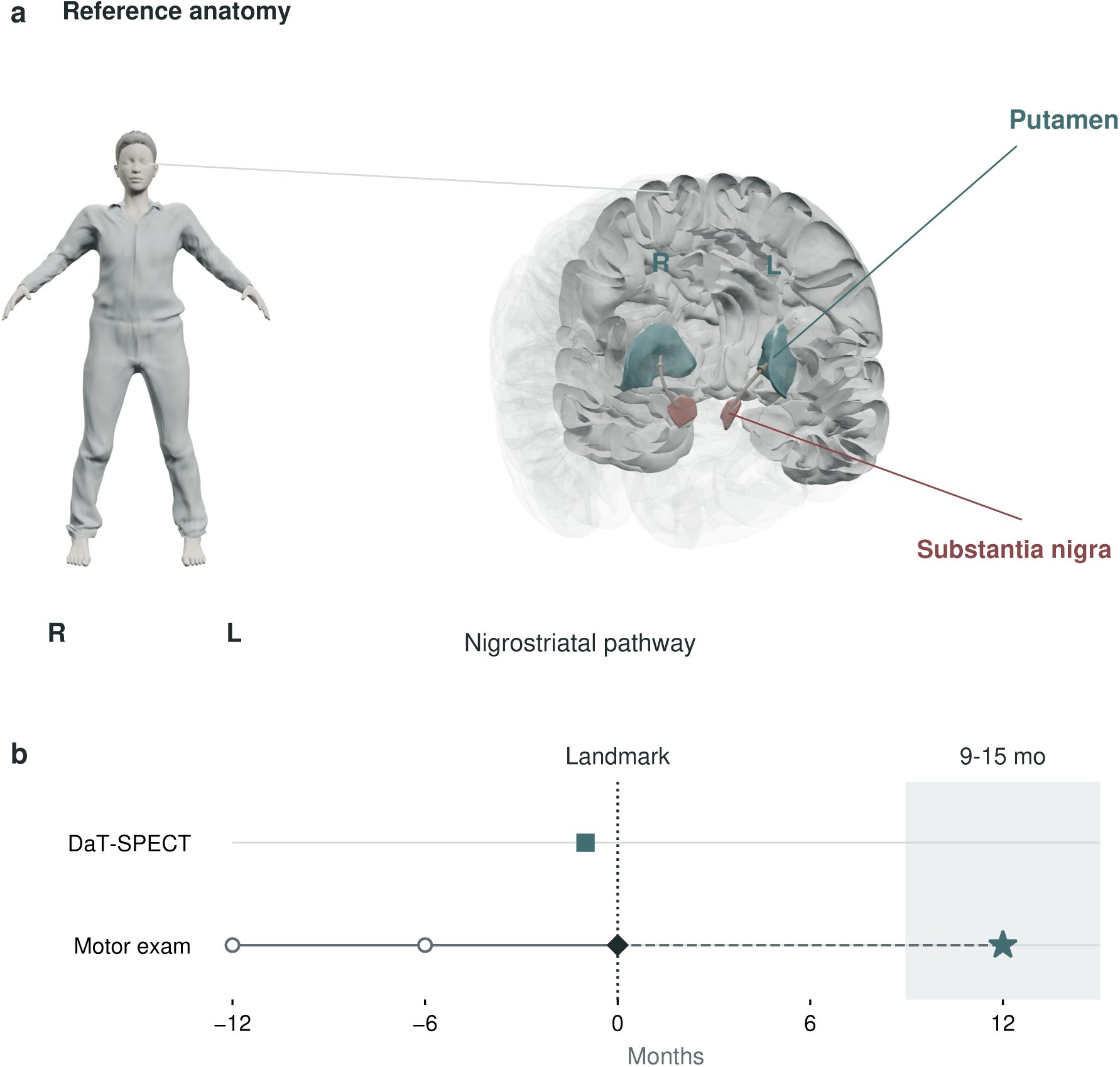
Anatomical interpretation and prospective evaluation answer different questions. a, Reference anatomy locates bilateral putamen (teal) and substantia nigra (rust). Nigrostriatal curves depict same-hemisphere projections schematically, not measured fibers. The atlas represents the substantia nigra region, not a separate pars compacta segmentation. Contralateral clinical expression is not a direct putamen-to-limb nerve; SBR is tracer binding, not a neuron count. b, Illustrative timeline: prior examinations (circles), scan (square), complete scan–examination landmark (diamond), and future examination (star). Pairing is within three months; all inputs must precede or coincide with the landmark. The shaded 9–15-month window defines the approximately 12-month endpoint. The analysis includes 1,602 PD participants with dated examinations, 1,439 with bilateral putamen SBR, and 1,204 eligible one-per-person landmarks. Evaluation is participant-held-out; MoCA, SAA and CSF context do not enter these forecasts. Reference human: MakeHuman/MPFB, CC0; brain: NIH 3D Human Reference Atlas (3DPX-020960), CC BY 4.0.

Before fitting a model, we made the intended use of each source explicit. The focused forecast uses current clinical state, demographic and treatment covariates, and signed DaT-SPECT. Cognition, SAA and CSF remain visible as dated biological context but do not enter this forecast. This distinction lets a reader see whether an apparent state is supported by contemporaneous directional evidence, a stale contextual measurement, or genuine disagreement between sources. Figure 2 connects the examined body, putamen and named nigrostriatal pathway, and places this anatomical interpretation inside the future-only design used for evaluation.

### 2.4 A direction-aware landmark asks which side rather than only how much

The landmark operationalizes a two-part dopaminergic state. Mean putamen SBR indexes overall bilateral DaT availability, with lower values reflecting greater deficit; signed asymmetry captures how that deficit is distributed across hemispheres. Patients can therefore match on mean SBR, total motor burden and even absolute asymmetry while occupying opposite spatial states. A near-symmetric record represents little directional imaging evidence, whereas an imaging–clinical conflict represents strong but internally inconsistent directional evidence. These are differences in the spatial expression and coherence of one mechanism family, not proof of different disease etiologies.

The PPMI data freeze contained 1,602 participants classified as PD with dated motor examinations. Of these, 1,439 had an analyzed DaT-SPECT record with valid left and right putamen specific-binding ratios (SBRs), and 1,204 had an eligible scan-clinical landmark plus a future motor examination at 12 months. The design selected the earliest valid landmark for each participant, so 1,204 is both the number of records and the number of independent people in the primary analysis. No participant contributed repeated opportunities to the primary prediction test.

At the landmark, mean age was 63.1 years (SD 9.6), 449 participants (37.3%) were recorded female, median diagnosis-derived disease duration was 0.67 years (interquartile range 0.31 to 1.58), and mean MDS-UPDRS Part III was 22.1 (SD 10.3). Age was unavailable for one participant. The median scan-clinical gap was zero months: 636 pairs occurred in the same month, 284 were separated by one month, 204 by two months and 80 by three months. Follow-up laterality was available for 1,204 participants at 12 months, 901 at 24 months and 682 at 36 months.

**Table 2:** Characteristics and analytical roles of the primary and independent replication cohorts. Values are mean (SD), median (interquartile range), or number (%). The cohorts were not pooled, and no between-cohort hypothesis test was performed.

| Characteristic | PPMI longitudinal cohort<br>(n=1,204) | S4 biological replication (n=56) |
| --- | --- | --- |
| Analytical role | Future 12-month laterality | Cross-sectional OFF-state replication |
| Age, years | 63.1 (9.6); 1 missing | 63.0 (8.2) |
| Recorded female | 449 (37.3%) | 19 (33.9%) |
| Disease duration, years | 0.67 (0.31-1.58) | 3.3 (1.1-5.8) |
| MDS-UPDRS Part III total | 22.1 (10.3) | 26.5 (11.8) |
| Scan-clinical separation | 0 months (0-1) | 17 days (8-42) |
| Motor examination state | OFF 131; ON 51; unknown/ambiguous 1,022 | OFF 56 |
| MoCA observed by landmark | 1,193 (99.1%) | Not used in replication |
| SAA observed by landmark | 616 (51.2%) | Not used in replication |
| Paired CSF alpha-synuclein and NfL | 381 (31.6%) | Not used in replication |
| LEDD history observed by landmark | 313 (26.0%) | Not used in replication |

We represented clinical laterality as the normalized right-body minus left-body burden across the same 11 paired MDS-UPDRS Part III domains. We represented imaging laterality as right putamen SBR minus left putamen SBR, divided by their sum. A positive imaging value therefore indicates relatively lower left putamen binding, whereas a positive clinical value indicates greater right-body burden. Positive correspondence is consequently compatible with the expected contralateral nigrostriatal relation. The primary outcome was the continuous clinical ratio at the future examination, not change in total MDS-UPDRS Part III and not a binary side label.

The landmark was the later month of the scan and its nearest clinical examination within three months. Each future outcome had to occur after that complete multimodal landmark. The 12-month examination was selected nearest 12 months within a 9- to 15-month window; the same rule was applied at 24 and 36 months. This ordering prevents a future clinical value from entering the baseline record and makes the temporal question explicit.

Medication-state information in the primary landmark was incomplete and is therefore a major boundary, not an implicit adjustment success. The baseline motor examination was marked OFF for 131 participants, ON for 51 and unknown or ambiguous for 1,022. Active LEDD history was observed at the landmark for 313 participants, of whom 303 had positive exposure. The primary models retained state indicators, LEDD value and an LEDD-availability indicator. A dated secondary landmark analysis then required the baseline and outcome to carry the same explicit OFF or ON label by construction.

### 2.5 Outcome-blind patient histories expose both resolution and uncertainty

Population summaries do not show what source-aware integration changes in one record. We therefore prespecified an outcome-blind visualization protocol before selecting any example. Eligibility required a same-month scan and motor examination, pre-landmark MoCA, definitive SAA, paired CSF alphasynuclein and NfL within 12 months, observed treatment context and all four held-out forecasts. Thirty-two participants met every requirement. No future value, future sign or forecast error entered eligibility, archetype assignment, ranking or tie-breaking.

Four records were selected to make different information states visible rather than to display four favorable outcomes. Cases A and B carried concordant, mirror-image putamen and body-side signs at the landmark; Case C had near-zero putamen asymmetry; Case D had opposing imaging and clinical signs. Figure 1 shows the two concordant mirror states at publication scale. Supplementary Figure 2 documents the prespecified selection process, and Supplementary Figures 3–6 retain the full Case A–D dossiers, including the near-symmetric and conflicting records. Across the displays, relative source times reveal whether cognition, SAA and CSF were current or stale; separate left-right values reveal anatomy that a mean or absolute asymmetry would erase; and the post-landmark trajectory reveals what happened only after selection.

The examples make patient difference tangible. Case A had left/right putamen SBR of 0.62/1.07 and paired left/right motor burden of 9/11; Case B had SBR of 1.26/0.56 and motor burden of 6/2. These are opposite concordant configurations, not a severity-matched pair. Case C’s almost symmetric SBR (0.58/0.57) coexisted with strongly asymmetric motor burden (1/11). Case D’s SBR (0.15/0.76) pointed toward greater right-body burden, but the examination showed the opposite (12/9). Keeping the measured sides visible distinguishes concordance, weak imaging direction and genuine cross-source conflict. None establishes a distinct molecular subtype or treatment pathway.

The four-model comparison makes the contribution, and its limits, inspectable (Table 3). Signed imaging moved Case B’s forecast closer to its observed left-sided predominance, but moved Case A’s forecast farther from the subsequently symmetric examination. In Case C, it reduced but did not resolve the overprediction of right-sided predominance. In Case D, it changed the predicted side toward the later observation but overestimated its magnitude; the magnitude-only forecast happened to be closer to that outcome. These are transparent prediction contrasts, not patient-level estimates of an imaging intervention’s effect. MoCA, SAA and CSF differences provide context for further investigation, not demonstrated explanations for these outcome differences.

**Table 3:** What directional imaging changes in each illustrative record. Values are held-out predictions or observed signed clinical laterality, not total motor scores. The four cases were selected without their future outcomes; they do not estimate average benefit.

| Case | Baseline | Clinical | + mean SBR | + magnitude | + signed SBR | Observed |
| --- | --- | --- | --- | --- | --- | --- |
| A | 0.100 | 0.106 | 0.107 | 0.125 | 0.249 | 0.000 |
| B | -0.500 | -0.322 | -0.320 | -0.286 | -0.472 | -0.500 |
| C | 0.833 | 0.647 | 0.651 | 0.630 | 0.538 | 0.000 |
| D | -0.143 | -0.073 | -0.067 | 0.006 | 0.333 | 0.143 |

Individual forecast uncertainty remained substantial. The cross-fitted 90% conformalized quantile intervals covered 95.5% of the 1,204-person cohort, but their median width was 1.504 on a signed-laterality scale bounded from -1 to 1. Only one of the two concordant mirror cases retained the imaging-indicated sign at 12 months. These were therefore not retrospectively curated success stories. They show a more useful distinction: anatomy and examination can resolve the current directional state while future individual laterality remains uncertain, and cross-source conflict can be preserved rather than averaged into a falsely precise state.

### 2.6 Signed dopaminergic anatomy reproduces across PPMI and S4

Before asking a longitudinal question, we tested whether the sign convention recovered the expected cross-sectional physiology. In PPMI, 1,367 PD participants had a valid putamen scan and clinical examination no more than three months apart. Signed putamen SBR asymmetry correlated with signed clinical laterality at r=0.676 (two-sided Pearson *P* = 1.56 × 10*^−^*^183^; 95% participant-bootstrap CI 0.649 to 0.702; Spearman r=0.692). The partial correlation remained 0.675 (0.646 to 0.702) after adjustment for total motor severity and scan-examination gap. Among 1,330 non-tied pairs, the lower putamen SBR was contralaterally compatible with the more affected body side in 1,092, or 82.1% (95% Wilson CI 80.0% to 84.1%).

S4 supplied an independent cohort and a stricter clinical state. Among 56 PD participants with analyzed DaT-SPECT and complete OFF-state paired motor scores, the same sign convention produced r=0.551 (0.363 to 0.701; Spearman r=0.590; two-sided Pearson *P* = 1.08 × 10*^−^*^5^). The association remained after adjustment for total motor score, scan-examination gap and S4 stage (partial r=0.562, 0.375 to 0.707). Leave-one-out correlations ranged from 0.525 to 0.614, and winsorization yielded r=0.554, arguing against dependence on one extreme observation. Forty-one of 54 non-tied pairs were directionally concordant, or 75.9% (63.1% to 85.4%). The PPMI and S4 signed correlations were not detectably different by a Fisher transformation (P=0.147).

Discarding sign substantially weakened the relation. Absolute putamen asymmetry correlated with absolute clinical asymmetry at r=0.206 in PPMI (two-sided Pearson *P* = 1.59 × 10*^−^*^14^), less than one third of the signed correlation. In S4, the magnitude-only correlation was r=0.058 (95% bootstrap CI -0.182 to 0.294), and its adjusted partial correlation was 0.059 (-0.220 to 0.341; twosided Pearson P=0.672). The cross-cohort result therefore does not merely state that more unequal striata accompany more unequal motor scores. It shows that retaining left-right direction preserves a reproducible contralateral relation (Figure 3).

**Figure 3:**
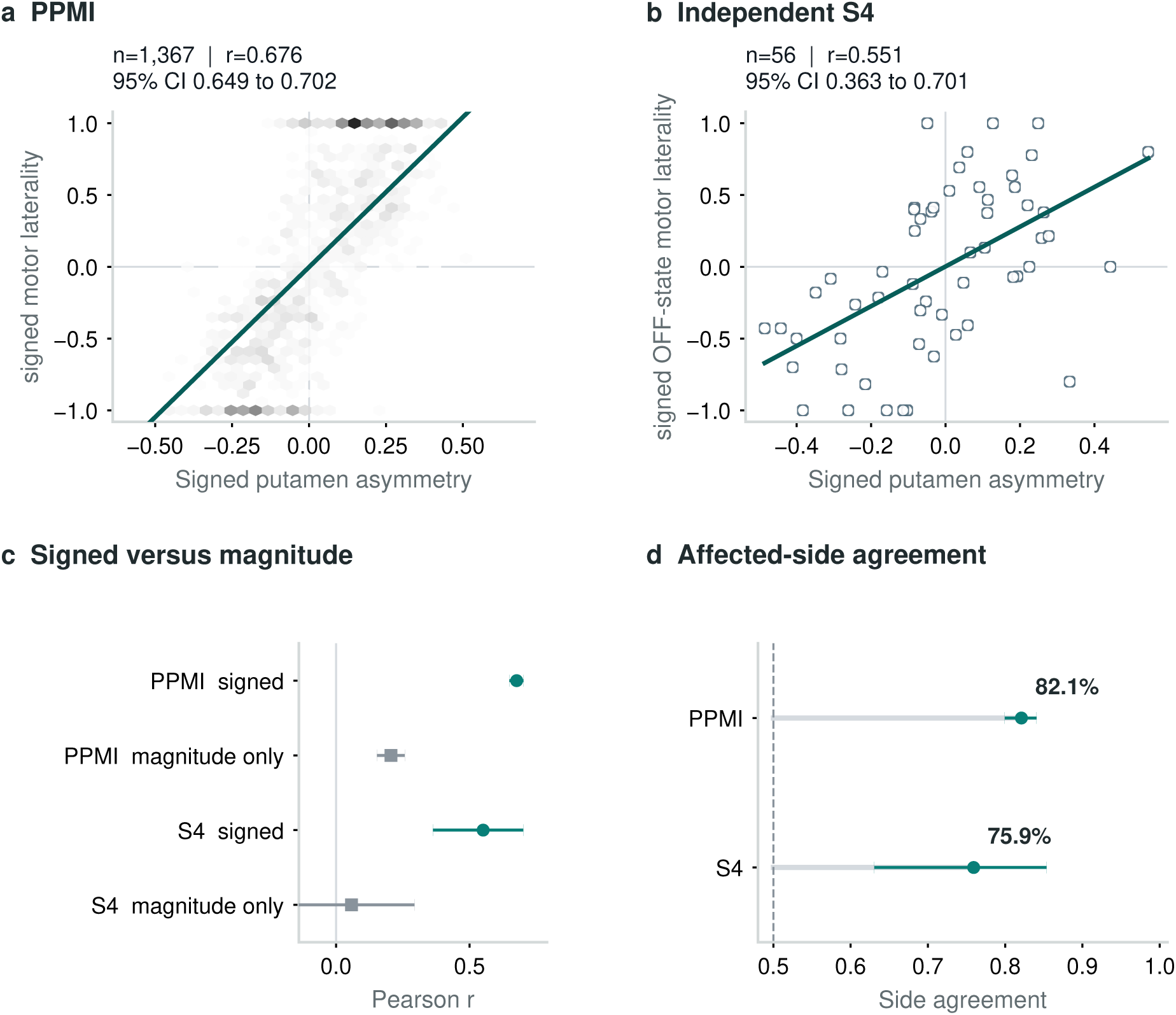
The signed contralateral association replicates across PPMI and S4 and exceeds magnitude-only encoding. a, Hexagonal participant density for 1,367 Parkinson’s Progression Markers Initiative (PPMI) scan-examination pairs separated by no more than three months. Positive imaging and clinical ratios indicate lower left putamen binding and greater right-body burden, respectively; negative ratios give the mirror configuration. The line is the unadjusted least-squares fit. b, Independent Systemic Synuclein Sampling Study (S4) OFF-state replication (n=56) using the same sign convention without cohort-specific sign selection. Open circles are individual participants and the line is the unadjusted least-squares fit. c, Pearson correlations with two-sided 95% participant-bootstrap intervals for signed (teal circles) and magnitude-only (gray squares) encodings; n=1,367 for PPMI and n=56 for S4. d, Contralaterally compatible affected-side agreement with two-sided 95% Wilson intervals among non-tied pairs (n=1,330 PPMI; n=54 S4); the dashed vertical line marks 50% agreement. Correlations and side agreement are two descriptions of the same signed relation, not independent confirmatory endpoints.

A secondary baseline-matched audit quantified how often similar non-directional summaries could conceal opposite states. Among 1,056 eligible primary-cohort participants, fixed calipers on total motor burden, mean SBR and absolute asymmetry yielded 365 non-overlapping pairs with opposite imaging signs; 281 pairs also had opposite clinical signs. These counts describe directional information discarded by unsigned summaries, not misclassification by the clinical model, which already includes clinical sign. Median within-pair differences in the saved future forecasts were 0.736 for clinical state and 0.823 for signed imaging. Larger separation alone does not establish greater accuracy; the full-cohort held-out comparisons below provide that test. Supplementary Section S7 reports the matching rules and the earlier null biomarker-profile comparison.

### 2.7 Signed putamen asymmetry adds modest 12-month information

The prespecified longitudinal model hierarchy was designed to isolate the value of direction. The clinical-state model contained baseline clinical laterality, total MDS-UPDRS Part III, age, recorded sex, diagnosis-derived duration, examination-state indicators, active LEDD and LEDD availability. The overall-DaT model added mean putamen SBR and scan-clinical gap. The magnitude-only model additionally used absolute putamen asymmetry. The direction-aware model instead added signed putamen asymmetry. Every model used the same five participant folds, median imputation with missingness indicators, robust scaling and ridge regression with alpha fixed at 10.

Clinical state was already a strong predictor of 12-month laterality because motor-side predominance is persistent. Participant-held-out *R*^2^ was 0.713, MAE was 0.234 laterality units and Spearman correlation was 0.853. Mean putamen SBR without direction did not improve that reference (*R*^2^ 0.712; MAE 0.234), nor did absolute asymmetry (*R*^2^ 0.712; MAE 0.233). The direction-aware model achieved *R*^2^ 0.730, MAE 0.227 and Spearman correlation 0.863.

Paired participant bootstrap comparisons localized the increment to sign. The direction-aware model improved *R*^2^ over clinical state by 0.016 (95% CI 0.007 to 0.026), over the overall-DaT model by 0.018 (0.008 to 0.027), and over magnitude-only imaging by 0.018 (0.009 to 0.028). Relative to the overall-DaT model, MAE fell by 0.0067 laterality units (-0.0111 to -0.0023). This is a stable but small increment. It should not be interpreted as a clinically important change on MDS-UPDRS Part III because the outcome is a normalized side-balance ratio, not the total motor score.

A separate model-refitting sensitivity, specified after the primary analysis, refitted the complete imputation, scaling and ridge pipeline in 1,000 participant bootstrap samples and evaluated each fit only on participants absent from that draw’s training sample. Relative to the overall-DaT model, the median out-of-bag Δ*R*^2^ was 0.0175 (2.5th to 97.5th percentiles 0.0015 to 0.0292), and 98.1% of draws favored signed direction. Median delta MAE was -0.0068 (-0.0129 to 0.0000), with 97.3% of draws favoring signed direction. These are refit-stability percentiles rather than an external-validation interval, but they show that the increment was not created only by one prespecified fold assignment. An adjusted association analysis reached the same conclusion without relying on out-of-fold prediction. After baseline laterality, total motor burden, mean putamen SBR, scan-clinical gap, age, recorded sex, disease duration, medication-state indicators and LEDD context were represented, a 0.1 increase in signed putamen asymmetry corresponded to a 0.0507 increase in 12-month clinical laterality (95% CI 0.0387 to 0.0626; partial r=0.251). On the unnormalized scale, the same 0.1 asymmetry corresponded to 0.757 additional right-minus-left paired motor points (0.600 to 0.918) after adjustment for the baseline right-minus-left difference and the same context. The direction was therefore not created solely by normalizing the clinical outcome.

The model did not improve the cruder question of which side would be worse. Future-side accuracy was 90.2% for clinical state and 89.8% for the direction-aware model; the paired difference was -0.3 percentage points (95% CI -1.4 to 0.7). The current clinical examination already identified the future dominant side for most participants. Imaging contributed modest continuous information about how strongly that side predominance remained expressed, not a better binary side decision.

**Table 4:** Clinical interpretation boundaries for the signed-laterality results.

| Result | Clinician-facing meaning | What must not be inferred |
| --- | --- | --- |
| PPMI/S4 signed correlation | DaT hemisphere and contralateral motor side align reproducibly | Not a new diagnostic test |
| Adjusted 12-month association | Direction carries future side-balance information | Not global motor progression |
| No binary-side improvement | The examination already identifies the dominant side for most people | Not a reason to order DaT-SPECT |
| Same-state association persists | Recorded OFF/ON matching does not erase the association | Not medication-invariant prediction |
| Molecular/cognitive tests fail FDR | No contextual modifier was supported | Not a molecular subtype |

### 2.8 Medication-state, onset-side and robust-outcome checks preserve the association

The adjusted signed-putamen association remained positive at the prespecified secondary horizons, although its magnitude declined. Separate horizon models estimated effects per 0.1 putamen asymmetry of 0.0370 at 24 months (95% CI 0.0228 to 0.0517; n=901) and 0.0242 at 36 months (0.0095 to 0.0388; n=682). An additional participant-clustered analysis retained every available horizon and resampled all observations from each participant together. Its corresponding effects were 0.0557 at 12 months (0.0446 to 0.0672), 0.0377 at 24 months (0.0256 to 0.0501) and 0.0132 at 36 months (0.0004 to 0.0260). The lower 36-month estimate reinforces attenuation rather than a stable long-range effect. Signed caudate asymmetry and the unnormalized right-minus-left outcomes showed compatible secondary patterns (Supplementary Information).

Association persistence did not imply stable incremental prediction. At 24 months, direction-aware *R*^2^ was 0.681 versus 0.673 for the overall-DaT model, but the paired Δ*R*^2^ interval crossed zero (0.008, 95% CI -0.001 to 0.018) and so did the MAE interval. At 36 months, the corresponding Δ*R*^2^ was 0.004 (-0.005 to 0.013). Attrition reduced the later-horizon samples, treatment and compensatory influences accumulated, and baseline clinical laterality remained a strong reference. We therefore treat 24- and 36-month results as support for a waning association, not as confirmed predictive gains.

Three harder evaluations tested whether the 12-month increment depended on pairing flexibility or a convenient split. In the 636 exact-month scan-clinical pairs, the adjusted effect per 0.1 putamen asymmetry was 0.0450 (95% CI 0.0307 to 0.0600; partial r=0.239). In a site-disjoint evaluation spanning 977 participants and 50 acquisition sites, direction-aware imaging improved *R*^2^ over the overall-DaT model by 0.0146 (0.0056 to 0.0238) and reduced MAE by 0.0062 (-0.0107 to -0.0017). No acquisition site appeared in both training and test folds.

For the strict-calendar test, training was restricted to 601 earlier entrants whose 12-month outcome was completed before 1 January 2020. The untouched test contained 582 participants first observed from 2020 onward. Direction-aware *R*^2^ was 0.712, compared with 0.693 for the overall-DaT model and 0.704 for clinical state. The directional increment over the overall-DaT model was supported (Δ*R*^2^ 0.0191, 95% CI 0.0066 to 0.0325; delta MAE -0.0076, -0.0135 to -0.0019). Its advantage over the clinical-only reference was uncertain (Δ*R*^2^ 0.0087, -0.0066 to 0.0248; delta MAE -0.0022, -0.0095 to 0.0051). Thus the calendar test confirms that preserving direction is better than adding non-directional imaging, but it does not establish that imaging is better than a well-measured clinical state under temporal deployment. The direction-aware calendar predictions also showed an intercept shift: their mean was 0.072 while the observed mean was 0.015, despite a calibration slope of 1.083. Preserved ranking therefore did not imply deployment-ready calibration.

The primary-cohort medication-state filter retained only 143 participants and was inconclusive (effect -0.0045 per 0.1, 95% CI -0.0363 to 0.0241). Because the primary landmark had been selected before applying that filter, a dated secondary analysis searched directly for the earliest scan and clinical pair with identical explicit follow-up state. This stricter design identified 688 participants: 532 OFF-to-OFF and 156 ON-to-ON. The adjusted association persisted at 0.0271 per 0.1 asymmetry (0.0132 to 0.0414; partial r=0.153). In contrast, direction-aware prediction improved *R*^2^ over the overall-DaT model by only 0.0063 (-0.0023 to 0.0148) and changed MAE by -0.0015 (-0.0053 to 0.0022). Holding medication state explicit therefore supports the anatomical association but not a stable incremental-prediction claim.

Historical side information provided a second triangulation. Among 1,160 participants with a lateralized onset side and non-zero imaging sign, signed putamen asymmetry agreed with the recorded onset side in 82.6% (95% Wilson CI 80.3% to 84.7%). Baseline and 12-month clinical signs agreed with onset side in 92.8% and 90.6%, respectively. Adding onset side to the adjusted design reduced the imaging effect to 0.0397 per 0.1 asymmetry (95% CI 0.0282 to 0.0518; n=1,169); additional adjustment for right, left or mixed handedness yielded 0.0398 (0.0281 to 0.0519; n=1,168). These related clinical measures are not independent replications, but they show that the imaging orientation follows the participant’s disease history and is not explained by recorded handedness.

All 1,204 primary landmarks had all 11 paired motor items observed on both sides at baseline and outcome; the complete-item estimate was consequently unchanged. Winsorizing raw baseline and future right-minus-left differences at the 1st and 99th percentiles retained an effect of 0.737 motor points per 0.1 asymmetry (0.581 to 0.897). Signed putamen asymmetry did not predict the non-lateralized sum of future gait and freezing items (coefficient -0.038, 95% CI -0.328 to 0.251). The primary direction-aware predictions were also well calibrated on their bounded endpoint (intercept -0.002, slope 1.016; median absolute error 0.185; no predictions outside -1 to 1). Together, these checks localize the finding to continuous side-specific expression rather than missing motor items, extreme raw differences or generic axial severity.

We additionally challenged the normalized endpoint with bounded-link models. Clipping the primary held-out linear predictions to [-1,1] changed no value and therefore reproduced the primary comparison. In an adjusted fractional-logit model of (laterality + 1)*/*2, each 0.1 increase in signed putamen asymmetry had an odds ratio of 1.126 for greater future right-sided balance (robust 95% CI 1.091 to 1.161). However, a participant-held-out ridge model fitted on the bounded logit scale did not preserve incremental prediction over the overall-DaT model (Δ*R*^2^ 0.0041, 95% CI -0.0057 to 0.0140; delta MAE 0.0038, -0.0007 to 0.0081). The signed association is therefore robust to a bounded mean model, whereas the small incremental linear forecast is link-dependent and is not supported as robust prognostic utility.

### 2.9 Imaging–clinical disagreement accompanies lower directional persistence

A signed coordinate also makes disagreement visible. Before inspecting this secondary result, we prespecified the hypothesis that greater baseline disagreement between putamen DaT-SPECT direction and clinical laterality would be associated with reduced future directional persistence and greater prediction uncertainty. Three definitions were fixed. *Directional disagreement* was the absolute difference between imaging and clinical laterality after each coordinate was centered at its median and divided by its interquartile range (IQR). A one-unit exposure contrast therefore means one IQR-standardized disagreement unit, not one conventional standard deviation. *Stable side* meant the same non-zero clinical sign at baseline and follow-up; a sensitivity excluded future ties rather than counting them as unstable. For display only, participants with absolute imaging asymmetry above 0.03 and matching signs were *concordant* ; opposing signs were *discordant* ; near-symmetric imaging and a zero clinical difference remained separate. Inference used the continuous disagreement coordinate and did not depend on that threshold. This is an evidence-quality coordinate, not a latent disease subtype or selective-prediction tool.

The pattern was common enough to test rather than illustrate. Of 1,204 participants, 893 (74.2%) were concordant, 163 (13.5%) discordant, 126 (10.5%) near-symmetric and 22 (1.8%) clinically indeterminate. The baseline dominant side remained dominant at 12 months in 93.8% of concordant participants, compared with 71.8% of discordant and 77.8% of near-symmetric participants. After adjustment for baseline clinical-laterality magnitude, absolute imaging asymmetry, mean putamen SBR, total MDS-UPDRS Part III, age, recorded sex, disease duration, scan–examination gap, acquisition site and medication-state indicators, each IQR-standardized increase in continuous disagreement had an odds ratio of 0.491 for retaining the baseline side (95% participant-bootstrap CI 0.297 to 0.711; n=1,178). Treating zero future side balance as indeterminate rather than unstable (OR=0.462, 0.257–0.703; n=1,153) and requiring exact-month scan–examination pairing (OR=0.425, 0.191–0.689; n=624) retained the association. In the much smaller explicit same-medication-state subset (n=138), the estimate was uninformative (OR=1.15, 95% CI spanning below and far above one); that analysis is non-confirmatory.

The categorical contrast was not an artifact of the 0.03 boundary. Discordant-versus-concordant stability odds ratios were 0.412 (95% CI 0.198 to 0.705) at a 0.02 imaging threshold, 0.335 (0.152 to 0.581) at 0.03 and 0.242 (0.092 to 0.428) at 0.05. Nor did the result depend on participants with only slight baseline clinical imbalance: continuous directional-disagreement odds ratios were 0.433 (0.239–0.667), 0.428 (0.223–0.683) and 0.417 (0.186–0.699) after requiring absolute clinical laterality above 0, 0.05 and 0.10, respectively. A participant-clustered analysis of 2,554 observations from 1,147 people across 12, 24 and 36 months also retained lower directional persistence (OR=0.629, 0.425–0.895 per IQR-standardized unit). Continuous disagreement increased absolute error from the primary out-of-fold direction-aware model by 0.0212 laterality units per IQR-standardized unit after adjustment (95% CI 0.0057 to 0.0377). A disagreement-informed conformal ranking changed risk–coverage area only trivially relative to clinical uncertainty (0.178 versus 0.179), so the result supports a readable evidence-quality coordinate rather than a new uncertainty model.

**Figure 4:**
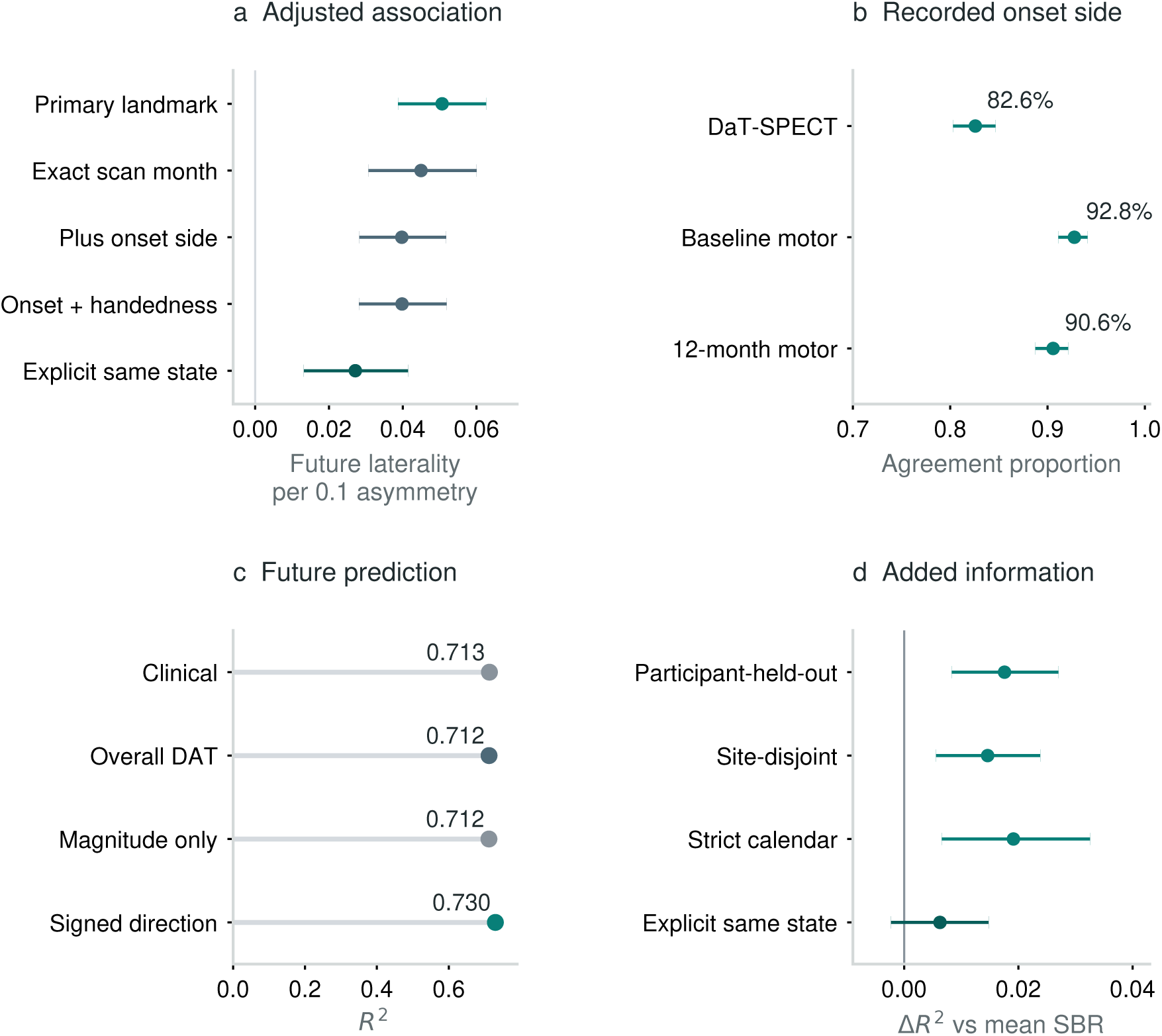
Signed anatomy tracks future laterality with modest predictive gain. a, Adjusted future clinical-laterality effects per 0.1 signed putamen asymmetry in the primary (n=1,204), exact-month (n=636), onset-adjusted (n=1,169), onset-and-handedness-adjusted (n=1,168) and explicit same-state (n=688) analyses. Points are estimates and horizontal lines are two-sided 95% participant-bootstrap intervals. b, Agreement with recorded side at onset for the DaT-SPECT sign (n=1,160) and baseline and 12-month clinical signs (n=1,146 each); horizontal lines are two-sided 95% Wilson intervals. c, Absolute participant-held-out 12-month *R*^2^ for clinical, overall-DaT, magnitude-only and signed-direction models in 1,204 participants; the axis begins at zero. d, Paired Δ*R*^2^ for signed direction versus the overall-DaT model under participant-held-out (n=1,204), site-disjoint (n=977), strict calendar (n=582 test participants) and retrospective explicit same-state (n=688) evaluation. Points are paired estimates and horizontal lines are two-sided 95% participant-bootstrap intervals; the vertical line is no increment. The adjusted association persists in the same-state sensitivity, whereas its incremental-prediction interval includes zero. The same-state cohort uses the recorded future examination state to define eligibility and is therefore a retrospective sensitivity analysis, not a prospective deployment test. Neither result demonstrates benefit for clinical decision-making.

Dated molecular and cognitive context was linked only from observations at or before the landmark and within the preceding 12 months, without imputing unmeasured assays. None of 11 prespecified comparisons passed Benjamini–Hochberg false-discovery control. SAA positivity was observed in 77.2% of 57 discordant records and 88.8% of 295 concordant records (unadjusted Fisher p=0.029; q=0.322); all continuous markers and MoCA were likewise null after correction. The screen therefore provides no molecular explanation for disagreement and argues against relabelling it as a biological subtype. The supported observation is narrower: two anatomically related sources conflict, and future side expression is correspondingly less persistent (Figure 5).

**Figure 5:**
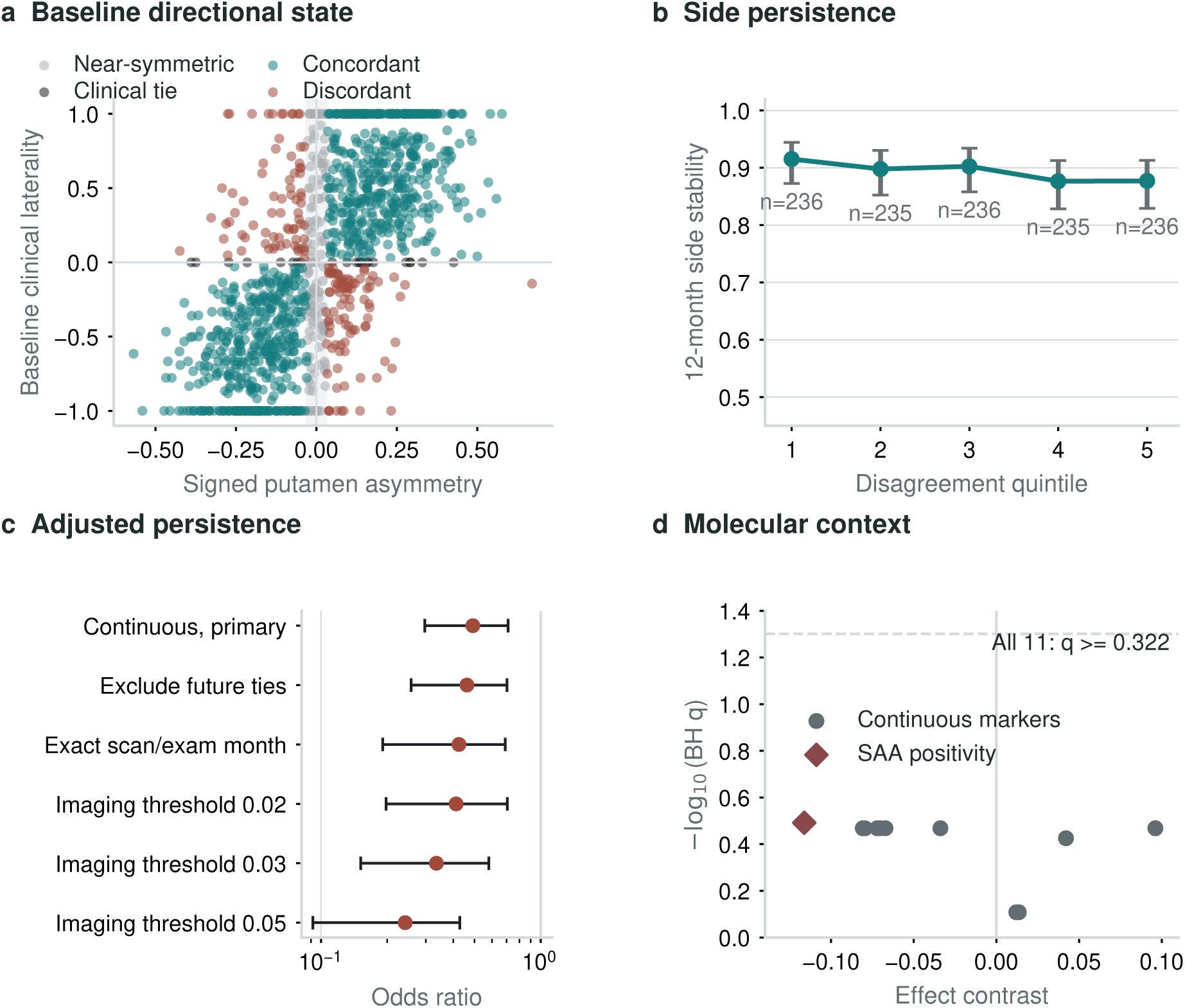
Directional disagreement makes lower persistence visible. a, Signed putamen asymmetry and clinical laterality in 1,204 PPMI participants: 893 concordant, 163 discordant, 126 near-symmetric and 22 clinically tied. The shaded band marks absolute imaging asymmetry at most 0.03; categories describe the data, whereas adjusted inference uses continuous disagreement. b, Twelve-month side persistence by disagreement quintile, with two-sided 95% Wilson intervals. c, Adjusted odds ratios for baseline-side persistence in the primary continuous, future-tie and exact-month analyses, then discordant-versus-concordant contrasts at imaging thresholds 0.02, 0.03 and 0.05. Points and lines are estimates and two-sided 95% participant-bootstrap intervals; the vertical line marks no association. The imprecise same-medication-state result remains in the supplement. d, Molecular/cognitive context linked within twelve preceding months. Gray circles show continuous-marker contrasts: Mann–Whitney probability of superiority for discordant versus concordant records, minus 0.5 (ties contribute half). The rust diamond instead shows the SAA-positive proportion difference, discordant minus concordant, not an odds ratio. Positive values indicate higher values or positivity in discordant records; the two effect measures are not interchangeable. The dashed line marks Benjamini–Hochberg (BH) q=0.05. No comparison survived false-discovery control; the screen does not establish a molecular subtype or cause.

## 3 Discussion

Preserving anatomical direction changed what multimodal data could credibly say. Signed putamen asymmetry reproduced the expected contralateral motor relation in two cohorts and tracked future clinical laterality after current laterality and total severity were represented. The association survived exact-month pairing, raw and winsorized outcomes, a bounded fractional-logit mean model, onset-side and handedness adjustment, participant-clustered repeated follow-up and an alternative 688-person landmark with identical explicit medication states. Prediction told a narrower story: a small 12-month linear increment over otherwise matched imaging models survived site and calendar separation, but it was uncertain under the bounded-link model and in the explicit same-state cohort. The result weakened at longer horizons, did not improve affected-side accuracy and did not transfer to a non-lateralized axial outcome. The supported claim is therefore specific: direction-preserving dopaminergic anatomy contains future-facing information about side-specific motor expression; robust incremental prediction and clinical utility are not established.

The biological significance is clearest in patient terms. Mean bilateral SBR indexes overall putamen DaT availability inversely: lower values indicate greater dopaminergic deficit. Signed asymmetry describes where that deficit is expressed. Two people can have similar mean SBR and total MDS-UPDRS Part III yet carry mirror-image hemisphere–body configurations, while a third can have near-symmetric imaging and a fourth can have imaging and examination pointing in opposite directions. A conventional point prediction may assign similar outcomes to these records. The direction-aware state additionally reveals the spatial mechanism coordinate and whether two anatomically linked sources agree. That explanatory distinction, rather than the small change in MAE, is the principal contribution.

The discordance analysis adds a second, complementary result. A direction-aware record can show not only which side is biologically compatible, but whether imaging and examination agree about that side. Discordance was associated with lower side persistence and greater future error after current imaging magnitude, mean putamen binding and clinical severity were represented. Yet adding disagreement to an uncertainty ranker barely changed risk-coverage area. This combination matters: cross-source conflict is a readable marker of a less stable evidence coordinate, but the current implementation is not a clinically useful abstention model. The entirely null pre-landmark molecular and cognitive screen further cautions against interpreting discordance as a discovered biological subtype.

The outcome-blind illustrative histories add an interpretive layer that aggregate correlations cannot supply. They show which sources were contemporaneous, which hemisphere and body side carried the sign, how the signed coordinate shifted a held-out forecast and where that forecast remained wrong or uncertain. Because eligibility and ranking never used the future outcome or forecast error, the records are not selected success stories. The cohort-level uncertainty analysis supplies the empirical boundary: cross-fitted 90% intervals covered 95.5% of participants but occupied a median 1.504 units of the two-unit outcome range. A source-aware representation can therefore resolve the anatomy of a current state while leaving an individual future state weakly resolved. These histories explain that distinction; they do not establish individual clinical utility.

This result is biologically plausible without being a causal mechanism claim. The putamen participates in the contralateral motor circuit, and lower regional dopamine-transporter availability is expected on the hemisphere opposite the more affected body side. Independent PET work has similarly related putamen DaT asymmetry to motor asymmetry [3], and the present PPMI-to-S4 replication shows that the relation survives a cohort change and an OFF-state clinical definition. The analysis does not show why one hemisphere became more affected, whether the asymmetry reflects alpha-synuclein propagation, differential vulnerability or compensation, or whether modifying DaT signal would alter motor laterality. Adjustment for recorded handedness did not remove the association, but handedness is not a randomized exposure and cannot resolve those mechanisms. The study identifies a directionally coherent in-vivo marker, not the cause of asymmetry.

The finding also clarifies an apparent disagreement in the literature. Cotogni and colleagues found no association between baseline asymmetry and later change in total motor, disability or non-motor burden [5]. Fiorenzato and colleagues reported hemisphere-associated differences in four-year motor and cognitive trajectories among strongly asymmetric, right-handed participants [4]. Our primary endpoint differs from both: it retains signed motor laterality as a continuous future state and conditions on the current signed state. The result does not imply that a more asymmetric participant will progress faster or slower. It says that the orientation of dopaminergic loss remains informative about the orientation and degree of later motor imbalance.

The magnitude-only control is central to that interpretation. Absolute asymmetry measures the degree of imbalance but makes mirror-image patients indistinguishable. It was weakly related to absolute motor asymmetry in PPMI, null in S4 and no better than mean bilateral putamen SBR for future signed laterality. The result is not an argument against unsigned biomarkers when the question concerns total severity. It is an argument that representations should retain the structure required by the target. In a lateralized biological system, sign is not a cosmetic encoding choice.

The high *R*^2^ of the clinical reference needs equally careful interpretation. Baseline laterality alone carries substantial information because the more affected side often remains more affected. A model that predicts the dominant side from the current examination is therefore already near 90% accurate. The imaging increment occurs on top of this strong persistence signal and changes MAE by less than one hundredth of the normalized laterality scale. That is scientifically informative because it isolates a reproducible anatomical contribution, but it is not evidence that ordering DaT-SPECT would improve a clinical decision. No decision-curve analysis, prospective intervention or clinically accepted threshold exists for this ratio.

The immediate clinical relevance is not that imaging identifies the more affected side better than a neurological examination; clinicians already observe laterality directly. The value is that multimodal records, longitudinal models and trial analyses need not erase the biological phenotype when they integrate sources. Retaining sign prevents opposite anatomical states from cancelling in a cohort average, supports side-aware monitoring and stratification in research, and exposes imaging–clinical conflict instead of converting it into false agreement. These are prerequisites for interpretable precision research, not evidence that one person should receive a scan, a drug or a prognosis in clinical care.

The temporal and site tests answer narrower questions than external validation. Site-disjoint folds reduce the opportunity to memorize acquisition-center patterns within PPMI. The strict-calendar split prevents later participants and their outcomes from influencing training. Both preserved the advantage of signed over non-directional imaging. However, under the calendar split the advantage over clinical state was uncertain, and neither design recreates a new health system, tracer workflow or independent longitudinal cohort. S4 validates the cross-sectional sign, not the PPMI longitudinal model. A diagnosis-linked external dataset with bilateral DaT measures, paired motor items and future examinations remains necessary before claiming longitudinal transport. Its analysis plan was specified before access to any new external outcome and prohibits cohort-specific sign reversal, refitting, recalibration and outcome-informed eligibility; that specification limits future analytical flexibility but is not itself external evidence.

Medication state remains an important measurement issue even after the larger same-state analysis. Reselecting the landmark around explicit OFF-to-OFF or ON-to-ON examinations retained 688 participants and supported the adjusted association, which makes the earlier 143-person null filter less concerning. Selection into that sensitivity was nevertheless strong. Compared with the 1,022 primary-cohort participants whose baseline state was unknown, the explicit same-state cohort had longer disease duration (2.70 versus 0.91 years; standardized mean difference 1.17), higher baseline MDS-UPDRS Part III (25.5 versus 21.7; standardized mean difference 0.35) and far more observed LEDD history (99.6% versus 13.0%; standardized mean difference 3.57). Its incremental-prediction interval crossed zero, OFF and ON groups were not randomized, and recorded state does not capture dose timing or long-duration dopaminergic response [18]. Future confirmation should prespecify practically defined OFF-state assessments, record dose timing and analyze treatment changes between landmark and outcome. The current result cannot be interpreted as medication-invariant biology or population-representative validation.

The explicit same-state cohort is also intrinsically retrospective: eligibility requires observing the medication-state label of the future examination. It therefore tests whether the association persists among comparable recorded states but cannot define a prospective deployment population. Conditioning on an observed future state may additionally select a non-representative subset, so this analysis narrows uncertainty without strengthening the primary predictive claim.

Other limitations follow from the source data. The study is observational and uses regional SBR summaries rather than harmonized raw-image reprocessing. The right-left ratios assume that regional labels and body-side items are recorded consistently across sites. Clinical laterality is a bounded ratio and can change when the denominator changes even if the raw side difference is stable; the unnormalized sensitivity reduced but did not remove this concern. Because directional persistence is defined relative to the baseline clinical sign, regression to the mean can contribute to apparent instability. Adjustment for baseline magnitude and the 0.05 and 0.10 clinical-margin sensitivities reduce, but cannot eliminate, that concern. Follow-up declined from 1,204 at 12 months to 682 at 36 months, so attenuation may reflect both biological convergence and selective observation. Handedness and onset side were available as recorded covariates rather than prospectively standardized mechanistic measures; motor subtype, interval treatment changes and compensatory cortical activity were not jointly modeled. The negative axial result supports specificity but cannot exclude an unmeasured common cause. Multiple secondary horizons and regions were interpreted as supportive rather than new primary tests.

The decisive next experiment is an independent cohort with idiopathic PD, quantitative bilateral putamen imaging, a baseline item-level motor examination within three months and the identical future side map 9–15 months later. The primary comparison should remain signed direction versus the same overall-DaT model without direction, with calibration, side agreement and continuous error reported separately. Raw-image reprocessing could then test whether posterior putamen, caudate or voxel-level patterns improve on regional SBR without sacrificing interpretability.

More broadly, this study offers a practical rule for mechanism-aware modeling: preserve the biological coordinate before adding model complexity. A multimodal state should retain source identity, anatomical direction, measurement time, missingness and treatment context long enough for each to be challenged. Only after a biological relation, a future prediction and a transport test agree should the state be carried toward treatment-response research. Here they do not all agree, and that disagreement is informative. Signed anatomy supports modest future side-balance information; global progression and treatment utility remain open problems.

In a future therapeutic digital twin, signed dopaminergic anatomy could serve as one observed state coordinate alongside molecular, cognitive, motor and treatment histories. Progressing from that state to intervention simulation or optimal control would require externally validated dynamics, causal treatment-response data, explicit dose and timing, calibrated success and failure probabilities, and prospective evidence that a simulated policy improves patient outcomes. The current study supports one biological coordinate and delineates its limits. It does not simulate therapy or optimize control.

## 4 Methods

### 4.1 Study design and data-use declaration

This was a retrospective secondary analysis of de-identified data collected by PPMI and S4. The PPMI protocol was approved by the institutional review board or independent ethics committee at each participating site, and all participants provided written informed consent [19, 20]. S4 study and recruitment materials were approved by the institutional review board or ethics committee at each site; participants provided written informed consent before study evaluations [17]. The parent studies were conducted in accordance with their approved protocols. No new participants were recruited and no new human measurements were collected for this analysis. Participant-level data were accessed under the applicable program agreements and were not redistributed. Empirical figures report aggregate analyses or outcome-blind, de-identified illustrative histories with arbitrary one-use labels, relative time and rounded derived values; no identifier, calendar date or participant scan is displayed.

The PPMI release used for this work was downloaded on 20 December 2025. The longitudinal signed-laterality protocol, model hierarchy, primary endpoint, bootstrap count, seed and decision rule were prespecified on 9 August 2026 before inspection of the longitudinal landmark result. Before the final 10,000-bootstrap analysis, a dated robustness protocol added the raw right-minus-left outcome, exact-month pairing and site-disjoint evaluation. Further protocols dated 12 and 19 August specified the alternative explicit same-state landmark; onset-side and handedness adjustment; complete-item, winsorized-outcome and bounded-link checks; participant-clustered repeated horizons; calibration summaries; medication-selection analysis; and the complete multiplicity audit. Because these extensions followed the primary analysis, they are reported as secondary and cannot replace or redefine the primary 12-month result. The dated protocols are included with the analysis code.

The study separates two levels of evidence. Cross-sectional PPMI and S4 analyses test whether the sign convention reproduces established physiology. The one-landmark PPMI analysis tests future side-specific association and prediction. Neither estimates global motor progression, a causal treatment effect, formal NSD-ISS stage, individual mechanism probability or clinical recommendation.

### 4.2 Prespecified longitudinal analysis

Before inspection of the longitudinal result, the primary analysis specified one earliest eligible landmark per PPMI participant, a clinical examination within three months of valid bilateral DaT-SPECT, normalized right-minus-left motor laterality at 12 months as the primary endpoint, putamen as the primary region and S4 as the independent cross-sectional replication cohort. Clinical, overall-DaT, magnitude-only and signed-direction models used the same imputation, scaling, ridge estimator and five participant-held-out folds. Primary intervals used 10,000 paired participant-bootstrap replicates.

The primary criterion required a positive adjusted 12-month putamen association, paired Δ*R*^2^ above zero and paired delta mean absolute error (MAE) below zero for signed direction relative to the otherwise matched overall-DaT model, with both prediction increments confirmed in the strict-calendar test. Independent cross-sectional S4 replication could not substitute for longitudinal confirmation. A landmark is the last month at which all information for one future prediction is available; source-time validity requires every predictor to precede or coincide with that landmark; and participant-held-out evaluation prevents any person from contributing to both model fitting and evaluation within a fold.

### 4.3 Parkinson’s disease population and source harmonization

PPMI PD status was taken from the dated clinical spine constructed from the release diagnosis fields. A participant was eligible for the longitudinal analysis if they had valid putamen measurements in the DaT-SPECT SBR analysis table, a clinical examination within three calendar months, and a future side-specific examination nearest 12 months within a three-month tolerance. Left and right putamen SBRs had to be non-negative with a positive sum. The same regional validity rule applied to caudate when available for secondary analysis; caudate availability was not an additional primary-cohort inclusion criterion. Records from controls, SWEDD and other non-PD classifications were excluded from the longitudinal disease analysis.

Clinical examination month was resolved from the examination date when available and the information date otherwise. Scan month came from the DaT-SPECT acquisition date. Event labels were standardized before joining sources. Age was joined at the participant-event level, recorded sex at the participant level, and disease duration was calculated from the earliest recorded PD diagnosis month. Active LEDD and whether LEDD history was observed were carried from the dated treatment spine. A missing exposure history was not coded as untreated.

The source modules below preserve each measurement’s scale, timing and analytical role. All PPMI modules came from the release downloaded on 20 December 2025. Clinical state, demographic/treatment covariates and DaT-SPECT entered the focused forecast. Cognition and molecular measurements were used in dated displays and exploratory context/profile comparisons, not as primary forecast inputs. No causal model was fitted. The reusable representation retains participant identity, measurement side, observation time and source availability before forming a landmark. It therefore preserves distinctions that a pooled, undated feature table cannot recover; it does not reconstruct unmeasured cellular or circuit activity.

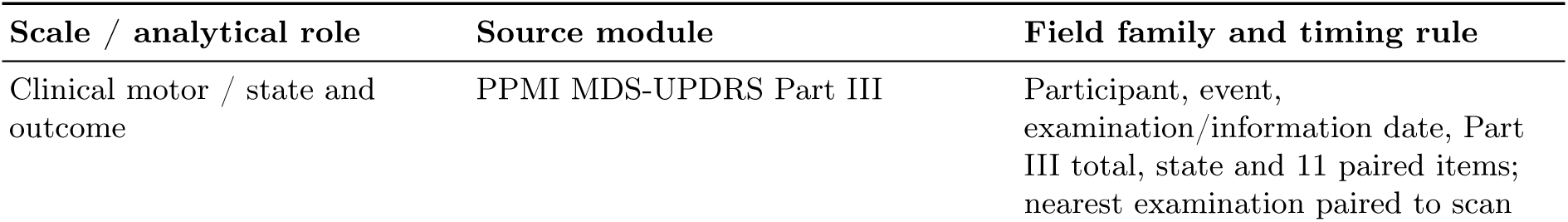
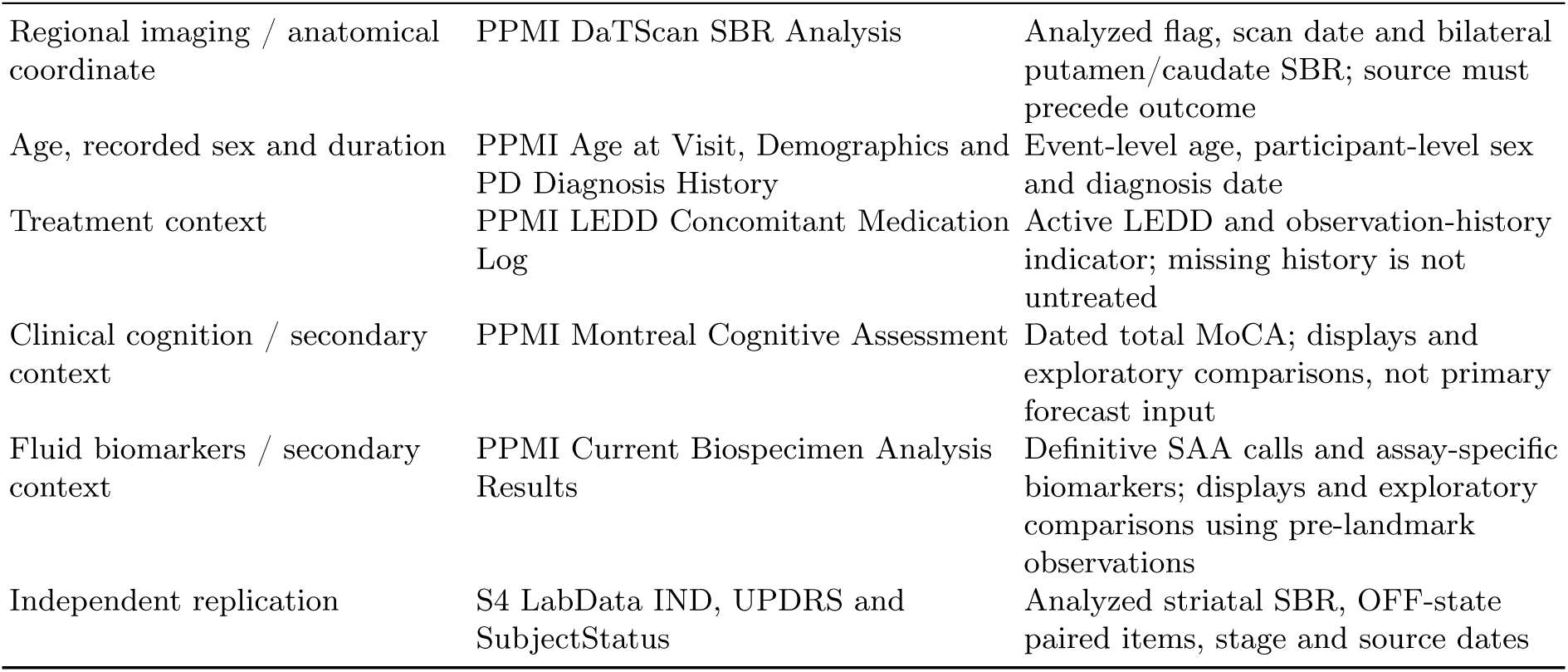

| Scale / analytical role | Source module | Field family and timing rule |
| --- | --- | --- |
| Clinical motor / state and outcome | PPMI MDS-UPDRS Part III | Participant, event, examination/information date, Part III total, state and 11 paired items; nearest examination paired to scan |
| Regional imaging / anatomical coordinate | PPMI DaTScan SBR Analysis | Analyzed flag, scan date and bilateral putamen/caudate SBR; source must precede outcome |
| Age, recorded sex and duration | PPMI Age at Visit, Demographics and PD Diagnosis History | Event-level age, participant-level sex and diagnosis date |
| Treatment context | PPMI LEDD Concomitant Medication Log | Active LEDD and observation-history indicator; missing history is not untreated |
| Clinical cognition / secondary context | PPMI Montreal Cognitive Assessment | Dated total MoCA; displays and exploratory comparisons, not primary forecast input |
| Fluid biomarkers / secondary context | PPMI Current Biospecimen Analysis Results | Definitive SAA calls and assay-specific biomarkers; displays and exploratory comparisons using pre-landmark observations |
| Independent replication | S4 LabData IND, UPDRS and SubjectStatus | Analyzed striatal SBR, OFF-state paired items, stage and source dates |

When duplicate clinical records remained in one participant-month and one examination state, numeric measures were collapsed by their median. If multiple states remained in the same month, explicit OFF was preferred, followed by explicit ON and then unknown or ambiguous state. This rule was specified without using the outcome value. Regional means and signed ratios were computed per scan row before their participant-month medians were taken; a median of ratios need not equal a ratio reconstructed from median left/right values.

### 4.4 Signed imaging and clinical laterality

For each region, signed imaging asymmetry was right SBR minus left SBR divided by right plus left SBR. Mean regional SBR was the arithmetic mean of right and left SBR and indexed overall DaT availability, with lower values indicating greater overall dopaminergic deficit. Magnitude-only asymmetry was the absolute value of the signed ratio. Positive putamen asymmetry therefore indicates relatively preserved right putamen binding and a larger deficit in the left putamen.

Clinical laterality used 11 paired MDS-UPDRS Part III domains [21]: upper- and lower-limb rigidity, finger tapping, hand movements, pronation-supination, toe tapping, leg agility, postural hand tremor, kinetic hand tremor, upper-limb rest tremor and lower-limb rest tremor. At least six items had to be observed on each side. Right and left item sums were computed separately. Signed clinical laterality was right-body minus left-body burden divided by their sum. A zero bilateral sum has no defined ratio and was represented as missing rather than assigned a side. Positive values indicate greater right-body burden. The unnormalized sensitivity used the raw right-body minus left-body difference.

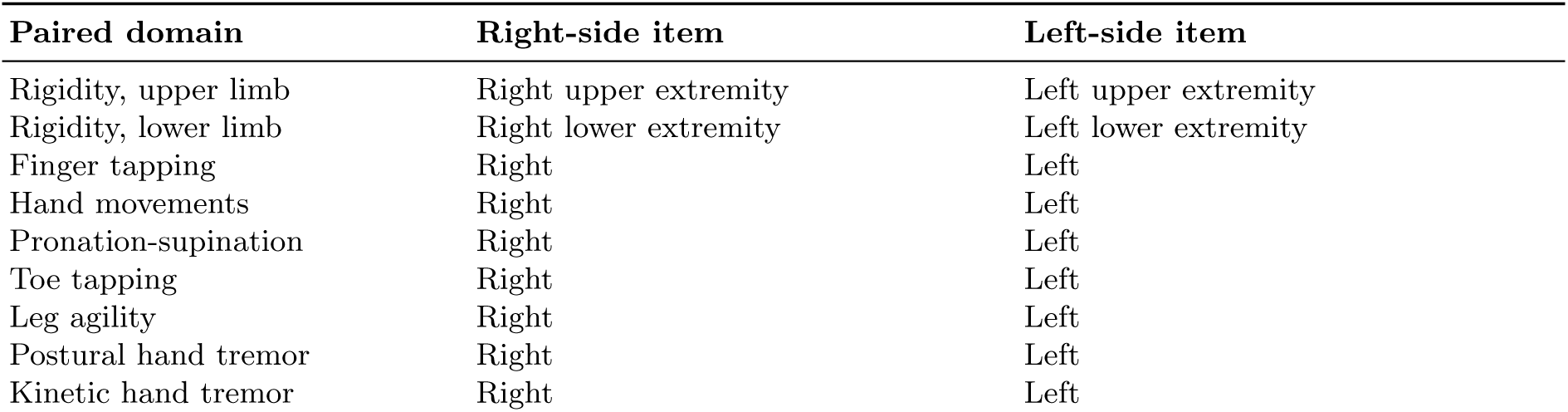
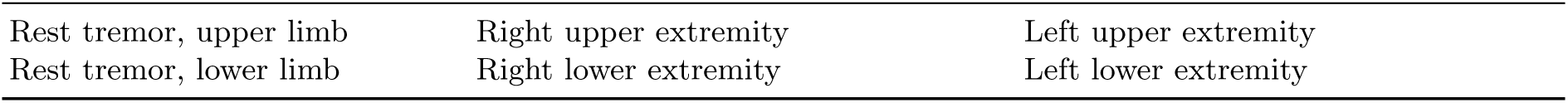

| Paired domain | Right-side item | Left-side item |
| --- | --- | --- |
| Rigidity, upper limb | Right upper extremity | Left upper extremity |
| Rigidity, lower limb | Right lower extremity | Left lower extremity |
| Finger tapping | Right | Left |
| Hand movements | Right | Left |
| Pronation-supination | Right | Left |
| Toe tapping | Right | Left |
| Leg agility | Right | Left |
| Postural hand tremor | Right | Left |
| Kinetic hand tremor | Right | Left |
| Rest tremor, upper limb | Right upper extremity | Left upper extremity |
| Rest tremor, lower limb | Right lower extremity | Left lower extremity |

Because striatal dopaminergic deficit is associated with contralateral body-side motor burden, a positive imaging ratio is expected to align with a positive clinical ratio: lower left putamen binding with greater right-body burden, or the mirror configuration for negative values. This clinical correspondence does not imply that nigrostriatal projections cross between hemispheres. The convention was held fixed across PPMI, S4, baseline and future analyses. No cohort-specific sign flip was permitted.

### 4.5 Cross-sectional biological replication

The PPMI cross-sectional set included one valid signed DaT-SPECT and clinical pair per PD participant with no more than three months separation. The nearest pairing rule was independent of the measured sign. We estimated Pearson and Spearman correlations, participant-bootstrap 95% intervals and partial correlations adjusted for total MDS-UPDRS Part III and scan-examination gap. The independent S4 analysis was restricted to PD stages 1 through 3 with an analyzed DaT-SPECT scan and sufficient OFF-state side-specific motor items. The same item map, imaging equation and sign convention were applied. The adjusted analysis included total motor score, scan-examination gap and S4 stage. Robustness checks reported leave-one-out correlation ranges and winsorized correlations. A Fisher transformation compared PPMI and S4 signed correlations. For a descriptive affected-side translation, exact zero imaging or clinical ratios were removed because no side could be assigned. Concordance required the lower-SBR putamen to be contralateral to the clinically more affected body side. Wilson intervals quantified the proportion. This proportion and the signed correlation describe the same biological relation and were not treated as two independent hypothesis tests. Absolute imaging-to-absolute clinical asymmetry was the magnitude-only negative control.

### 4.6 Longitudinal landmark

Each PPMI participant contributed at most one landmark. Candidate DaT-SPECT scans were ordered chronologically. For each scan, the nearest clinical examination within three months was selected, with timing preferred before examination state. The landmark month was the later of the scan and clinical months. The earliest pair with a valid 12-month future examination was retained. This ensured that both imaging and baseline clinical information existed before follow-up.

Future records had to occur after the landmark. For each horizon, we first minimized absolute distance to 12, 24 or 36 months within a three-month tolerance. Ties then preferred the baseline examination state, followed by OFF state and chronological order. The primary endpoint was 12-month signed clinical laterality. Twenty-four and 36 months were secondary. The non-lateralized sum of gait and freezing at 12 months was the negative-control outcome.

### 4.7 Outcome-blind patient histories and predictive uncertainty

The illustrative-history population was nested inside the primary 12-month cohort. Eligibility required zero-month scan-examination separation; a MoCA score, definitive positive or negative SAA call, and paired CSF alpha-synuclein and NfL observation no more than 12 months old and no later than the landmark; an explicit OFF/ON state or time-valid LEDD history; and complete out-of-fold predictions from the four prespecified model families. Thirty-two participants met these conditions. Four interpretable records were chosen without using the participant’s future value, future sign, forecast error or later trajectory. Case A required positive putamen asymmetry above 0.05 and concordant positive clinical laterality; Case B used the negative mirror definition; Case C minimized absolute putamen asymmetry within a maximum of 0.03; and Case D required putamen asymmetry of at least 0.05 in magnitude with an opposing clinical sign. Directional cases were ranked by stronger baseline anatomy, narrower cross-fitted uncertainty, lower disagreement among the four held-out model families and fresher source measurements. The near-symmetric and conflict cases used the corresponding prespecified low-direction and high-conflict rankings. Remaining ties were resolved deterministically without outcome information.

Patient-specific uncertainty used five separately generated outer participant folds (seed 20260824), not the primary ridge fold assignment. Within each outer training set, 20% of participants were reserved for calibration. Median imputation with missingness indicators was fitted only to the remaining participants. Gradient-boosted 0.05, 0.50 and 0.95 quantile regressors used the direction-aware feature set, 250 trees, learning rate 0.03, maximum depth two, minimum leaf size 20 and full-sample boosting. Lower and upper predictions were ordered before calibration. For calibration participant *i*, with ordered bounds *l_i_, u_i_* and outcome *y_i_*, the conformalized quantile regression (CQR) score was

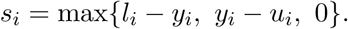

With *n*_cal_ calibration participants, the adjustment was the empirical higher quantile of these scores at probability min{1, ⌈0.90(*n*_cal_ + 1)⌉*/n*_cal_}. The adjustment was subtracted from each ordered test lower bound and added to its upper bound; both were then clipped to the valid [-1,1] outcome range. Nominal marginal coverage was 90%. Each interval is displayed with that quantile model’s own median, not with a ridge prediction. This exploratory analysis communicates uncertainty without replacing the primary comparison. Observed-history curves use the primary analysis’s examination-state collapse, with exact agreement checked against the saved baseline and selected future endpoint.

For display, CSF alpha-synuclein and NfL were converted to percentile ranks within the strict eligible visualization set because assay values were not treated as clinical reference thresholds. Calendar dates and participant identifiers were suppressed, source times were expressed relative to the landmark and displayed values were rounded. The internal identifier map and row-level derived tables remain controlled. Case-level and aggregate figures are governed by the same program data-use and publication requirements.

Reference brain anatomy was rendered reproducibly from the NIH 3D Human Reference Atlas “Brain, Male” mesh (3DPX-020960), based on the Allen Human Reference Atlas–3D, 2020 (version 1.0.0; RRID:SCR_017764) [22] and distributed under CC BY 4.0. Reference human surface and clothing meshes came from MakeHuman/MPFB’s CC0 assets. The same neutral reference pose was used across cases; body appearance, shape and motion were not reconstructed from participant data. Three.js supplied deterministic 3D renders, with vector labels and plots composed in Matplotlib. Cortical clipping exposes named putamen and substantia nigra meshes; pathway curves are schematic, not tractography. All four displayed records had one source examination at the selected landmark, and the 11 paired item sums exactly reproduced the frozen side totals. Figures and dossiers retain saved measurements and predictions without refitting or reselection. No generative image model was used.

### 4.8 Descriptive audit of information lost by unsigned summaries

This secondary display audit was specified on 11 September 2026 after the primary prediction analysis. It used the fixed 1,204-person primary cohort and excluded missing matching variables, absolute putamen asymmetry no greater than 0.03 and zero baseline clinical laterality. Candidate pairs had opposite imaging signs and absolute differences no greater than three motor-total points, 0.10 mean SBR units and 0.05 absolute-asymmetry units. Edges were ordered by the sum of squared caliper-scaled differences, with SHA-256 tie-breaking; greedy selection prevented participant reuse. Clinical-sign concordance, future values and forecasts did not rank pairs. After matching, we counted opposing clinical signs and compared the four frozen model predictions within pairs. These descriptive separations are not error rates, causal effects or an additional validation cohort. A separate earlier matched-profile analysis, including its null result and coverage sensitivities, is documented in Supplementary Section S7.

A separate earlier analysis tested secondary-profile differences among clinically and imaging-similar participants. Baseline matching used signed clinical laterality, total motor burden, mean and signed putamen SBR, age, disease duration, LEDD and LEDD availability. Variables were median-centered and interquartile-range scaled, with standard-deviation fallback; missing LEDD was median-imputed. Greedy non-overlapping Euclidean matching used the 75th percentile of nearest-neighbor distances as its caliper, with a 50th-percentile sensitivity. Secondary profiles and future outcome values did not form or rank pairs. Profile distance used robust-scaled MoCA, SAA, CSF alpha-synuclein and NfL, with prespecified reduced-coverage comparisons; future divergence was absolute paired 12-month laterality difference. Each reported run used 50,000 pair-bootstrap draws for percentile intervals and 50,000 permutations for two-sided tests. A standardized linear model also adjusted for residual baseline matching distance. These exploratory comparisons were not used to select the displayed cases.

### 4.9 Model hierarchy and internal evaluation

The clinical-state feature set comprised baseline signed clinical laterality, MDS-UPDRS Part III total, age, recorded sex code, diagnosis-derived disease duration, OFF and ON indicators, active LEDD and LEDD availability. The overall-DaT model added mean putamen SBR and absolute scan-clinical separation in months. The magnitude-only model further added absolute putamen asymmetry. The direction-aware model replaced absolute asymmetry with signed putamen asymmetry.

All models used a scikit-learn pipeline with median imputation, missingness indicators, robust scaling and ridge regression with alpha 10. Five shuffled participant folds were generated with seed 20260809 plus the horizon. Because each participant had one landmark, every prediction was participant held out. Feature definitions, estimator and hyperparameters were fixed; there was no outcome-driven feature selection, architecture search or tuning on held-out predictions.

We reported out-of-fold *R*^2^, MAE, RMSE, Spearman correlation, affected-side accuracy and affected-side area under the receiver-operating-characteristic curve among non-tied outcomes. Additional diagnostic summaries included median absolute error, calibration intercept and slope, prediction range and the fraction outside the physical interval from -1 to 1. No recalibration was applied. The primary comparisons subtracted the clinical, overall-DaT or magnitude-only metric from the direction-aware metric on the same participants. Ten thousand paired participant bootstrap samples produced 95% percentile intervals for Δ*R*^2^, delta MAE and delta side accuracy. Positive Δ*R*^2^ and negative delta MAE favor signed direction.

The bounded-outcome addendum separated association robustness from prediction robustness. First, primary held-out linear predictions were clipped to the physical interval [-1,1]; no prediction required clipping. Second, for each participant fold, the transformed training outcome (*y* + 1)*/*2 was clipped to [0.005,0.995], mapped through the logit, fitted with the unchanged ridge pipeline and mapped back to [-1,1] for held-out evaluation. Third, a fractional-logit mean model used (*y* + 1)*/*2 on [0,1], the complete adjusted association design and a logit link; a heteroskedasticity-robust sandwich covariance quantified the signed-asymmetry effect. No bounded model was recalibrated after validation.

To assess uncertainty from refitting rather than only resampling fixed predictions, a secondary out-of-bag bootstrap drew 1,204 participants with replacement 1,000 times. For every draw, median imputation, missingness indicators, robust scaling and all four ridge models were fitted anew on the bootstrap sample and evaluated only on participants absent from training. Hyperparameters and feature definitions remained fixed. We report the median, 2.5th and 97.5th percentiles and the fraction of draws favoring signed direction. Because out-of-bag sample composition varies and the procedure is not an independent cohort, these ranges are interpreted as model-refitting stability, not transport validation.

The prespecified decision rule required four findings: a positive adjusted 12-month putamen interval, direction-aware Δ*R*^2^ above zero relative to the overall-DaT model, direction-aware delta MAE below zero relative to that model, and confirmation of both increments in the strict-calendar test. Cross-sectional S4 replication could not substitute for the longitudinal criterion. Later horizons and caudate could support but not replace the primary claim.

### 4.10 Adjusted association and robustness analyses

We estimated the signed-putamen coefficient in an ordinary least-squares model containing baseline laterality, baseline total motor burden, age, recorded sex, disease duration, OFF and ON indicators, active LEDD, LEDD availability, mean putamen SBR and scan-clinical gap. Continuous missing covariates were median imputed. The coefficient was scaled to a 0.1 increase in signed asymmetry. Participant bootstrap intervals used 10,000 draws. Partial correlation was calculated by residualizing both signed putamen asymmetry and future laterality against the remaining design matrix.

Prespecified secondary analyses substituted signed caudate asymmetry and evaluated 24- and 36-month outcomes. The explicit-state sensitivity required both baseline and follow-up examinations to be identically marked OFF or ON; two unknown states did not count as a match. The negative control replaced future signed laterality with the non-lateralized gait-plus-freezing sum.

The dated robustness protocol repeated the adjusted analysis among same-month scan-clinical pairs and replaced the normalized outcome with the raw future right-minus-left paired-item difference. The raw-difference model controlled its baseline value, total motor burden and the same contextual covariates. These checks could narrow but not supersede the primary endpoint.

The 12 August secondary protocol rebuilt the landmark by requiring explicit OFF or ON examinations within three months of DaT-SPECT and an identically labelled 12-month outcome. Candidate scans were ordered chronologically; candidate baseline examinations were ordered by absolute scan gap, OFF before ON and date. The earliest pair satisfying the state-matched outcome criterion was retained. The same feature sets, estimator, folds and comparisons were repeated in this alternative cohort.

Side predominantly affected at onset was encoded left=-1 and right=+1 to match the clinical-laterality sign; symmetric and unknown onset categories were excluded from directional concordance. Wilson intervals summarized agreement of onset side with imaging, baseline clinical and 12-month clinical signs. Adjusted models then added onset side and recorded handedness indicators. A complete-item sensitivity required all 11 paired items on both sides at baseline and outcome. The raw right-minus-left analysis was repeated after 1st/99th-percentile winsorization. A pooled 12-, 24- and 36-month model included horizon indicators and signed-asymmetry-by-horizon interactions; participant-bootstrap samples kept all observations from each selected participant together.

### 4.11 Site-disjoint and strict-calendar evaluation

For site-disjoint evaluation, each participant was required to map to one acquisition site. Fifty represented sites were sorted by participant count and assigned greedily to five folds without using outcomes. All participants from one site remained in the same test fold. The model pipeline and feature sets were unchanged.

The strict-calendar evaluation used 1 January 2020 as the boundary. Training required an earlier first PD motor observation and a 12-month outcome completed before the boundary. Test participants were first observed from 2020 onward. Imputation, scaling and ridge fitting used only the 601-person training set; no recalibration or outcome-based selection was performed in the 582-person test set. Paired test-participant bootstraps quantified the directional increment.

### 4.12 Directional-disagreement analysis

The discordance addendum was locked on 17 August 2026 before its outcome-facing analysis. Near-symmetric imaging was defined as absolute putamen asymmetry no greater than 0.03. Above that threshold, a non-zero clinical laterality sign matching the imaging sign was concordant and an opposing sign was discordant; a zero clinical sign was clinically indeterminate. Thresholds 0.02 and 0.05 were prespecified sensitivities. The primary continuous exposure was the absolute difference between imaging and clinical laterality after each was centered by its median and divided by its interquartile range.

The primary binary endpoint indicated whether the sign of 12-month clinical laterality matched the baseline clinical sign; zero future laterality counted as unstable, with exclusion of future ties as a sensitivity. Adjusted logistic models included baseline laterality magnitude, absolute putamen asymmetry, mean putamen SBR, baseline total MDS-UPDRS Part III, age, recorded sex, diagnosis-derived disease duration, scan-examination gap, acquisition site, and OFF/ON indicators. Each bootstrap draw refitted imputation, scaling, site encoding and the adjusted model. Secondary adjusted linear models likewise refitted their complete design and tested aligned future laterality and absolute error from the primary out-of-fold direction-aware prediction.

For the uncertainty check, each held-out participant fold used three other folds to fit a ridge model of log absolute error and a fourth participant-disjoint fold for 90% conformal calibration. Clinical uncertainty used baseline clinical state and context; the disagreement-informed model additionally used mean and absolute putamen SBR, scan age and continuous disagreement. Risk was MAE after retaining predictions from lowest to highest estimated uncertainty.

Molecular observations, MoCA and definitive SAA calls were linked only when observed on or before the landmark and no more than 12 months earlier. Continuous markers were compared between concordant and discordant records using two-sided Mann–Whitney tests; SAA used Fisher’s exact test. CSF alpha-synuclein, amyloid-beta42, p-tau, t-tau, p-tau217, GCase, GFAP, NfL, NfL average, MoCA and SAA formed one Benjamini–Hochberg family. No missing assay was imputed for this screen. These analyses supplied biological context and could not establish a mechanism claim.

Clinical-sign margins of 0, 0.05 and 0.10 were prespecified to distinguish reproducible directional conflict from unstable signs near zero. At each margin we repeated the continuous standardized-disagreement analysis, the raw absolute imaging–clinical gap and the categorical sign-discordance contrast. A pooled 12-, 24- and 36-month analysis retained all eligible horizons and resampled complete participant trajectories; horizon indicators prevented different follow-up times from being treated as exchangeable rows.

### 4.13 Statistics and reproducibility

All intervals and tests are two-sided, and 95% intervals correspond to a nominal alpha level of 0.05. The 12-month putamen endpoint and direction-aware versus overall-DaT comparison were primary; caudate, later horizons, state matching, raw difference, exact-month pairing, site separation and the axial outcome were sensitivity or secondary analyses. No multiplicity correction was applied to these supportive analyses, so they were not interpreted as independent confirmatory findings. We emphasize effect estimates and intervals rather than a dichotomous P-value threshold. Exact Pearson and Spearman P values used their standard two-sided null distributions; the cross-cohort correlation comparison used a two-sided Fisher r-to-z test. Wilson score intervals were used for affected-side proportions. Bootstrap analyses report percentile intervals and, where generated, the proportion of bootstrap estimates above zero rather than converting resampling tails into parametric P values.

Cross-sectional robustness used 50,000 participant-bootstrap draws from the primary biological analysis; the prespecified longitudinal analysis used 10,000 draws and seed 20260809. The 12 August robustness analyses used 10,000 draws and seed 20260812. The directional-disagreement protocol used 5,000 participant-bootstrap draws and seed 20260817, with its three imaging thresholds, three clinical margins and multiple-testing family prespecified before outcome analysis. Disagreement inference refitted the complete adjusted model within every participant-bootstrap draw. Paired prediction-performance bootstraps resampled participants while holding their already fitted out-of-fold predictions fixed; those intervals quantify test-sample variation conditional on the prespecified folds and fitted predictions. The separate 1,000-draw out-of-bag analysis refitted the entire prediction pipeline to challenge that limitation; its percentile ranges are robustness ranges rather than external-validation confidence intervals.

Analyses used Python 3.11, pandas, NumPy, SciPy, scikit-learn and matplotlib. Cohort construction, split assignment, aggregate metrics and figure inputs were generated programmatically. Checks verified the sign equations, physical ratio bounds, one-landmark rule, future-only outcomes, scan-clinical windows, participant and site separation, model hierarchy and primary comparisons. Reporting was informed by TRIPOD+AI and STROBE principles [23, 24], while recognizing that the primary contribution is a prognostic association and representation audit rather than a deployable clinical prediction model. The Supplementary Information reports the complete secondary numerical record, patient-selection audit, bounded-outcome sensitivity and formal multiplicity family.

### 4.14 Use of AI-assisted tools

AI-assisted tools, including OpenAI Codex, were used only for manuscript revision, code development and debugging. The authors reviewed and verified all outputs and retain full responsibility for the manuscript, analyses and code.

## 5 Extended Anatomical and Patient Walkthroughs

### 5.1 Three numbers that should not be collapsed into one severity label

A bilateral scan provides an overall binding level, a degree of imbalance and a direction. For left/right putamen SBR *L_D_, R_D_*, these are

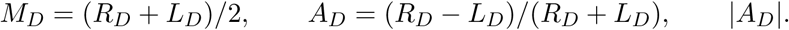

For paired motor sums *L_C_, R_C_*, the separate clinical coordinate is

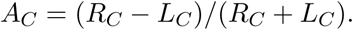

All ratios require a positive denominator. The motor total additionally contains unpaired findings and is not equal to *R_C_* + *L_C_*. Thus neither |*A_D_*| nor |*A_C_*| is a whole-disease severity score. The sign convention is fixed before comparisons: positive imaging asymmetry means relatively lower left binding, whereas positive motor laterality means greater right-body burden. Agreement has contralateral clinical meaning; it does not mean that nigrostriatal projections cross the midline.

Removing a sign is an irreversible representation choice. |*A_D_*| is unchanged if the two hemispheres are exchanged. A model receiving only the mean and magnitude cannot reconstruct the lost direction from those imaging features. This is why the primary comparison uses otherwise matched models rather than a larger model with many additional measurements.

### 5.2 Case A: current anatomical agreement does not guarantee persistence

Case A’s left/right putamen SBRs are 0.62/1.07. Their mean is 0.845, while (1.07 − 0.62)*/*(1.07 + 0.62) = 0.2663 identifies lower left binding. Left/right paired motor sums 9/11 yield (11−9)*/*(11+9) = 0.10, consistent with a relatively greater right-body burden. The actual right/left arm-rigidity scores are 2/1 and upper-limb rest-tremor scores are 1/0. Other items do not uniformly favor the right: the full paired profile is retained in the supplement. The clinical description is therefore right-predominant findings with rigidity and rest tremor, not a formal tremor-dominant subtype.

Figure 1a locates those measurements; panel c shows what happened later. At the selected future examination the clinical ratio was zero. The clinical-only forecast was 0.106, whereas the direction-aware forecast was 0.249. Keeping direction worsened this individual’s error despite the coherent initial anatomical state. The separate CQR median was 0.170 with a 90% interval approximately [-1.000,0.549]. That interval belongs to the quantile estimator and cannot be attached to the ridge prediction. The figure is informative precisely because it does not hide the failed individual forecast.

### 5.3 Case B: the opposite configuration with a better, but still uncertain, forecast

Case B’s left/right putamen SBRs are 1.26/0.56. Mean binding is 0.910 and signed asymmetry is (0.56 − 1.26)*/*(0.56 + 1.26) = −0.3846. Left/right motor sums 6/2 give laterality -0.50. Left-sided finger tapping, hand movements, pronation/supination and arm rigidity each score one while the corresponding right items score zero; toe tapping and leg agility score one on both sides. All paired limb-tremor items are zero. These findings support a left-predominant bradykinetic and rigid description without assigning an adjudicated motor subtype.

The clinical-only prediction was -0.322, the direction-aware prediction -0.472, and the observed future laterality -0.500. Imaging direction improved the point forecast for this case. The separate CQR median was -0.443 with a 90% interval approximately [-1.000,-0.089]. Its interval is narrower than Case A’s but still spans a substantial fraction of the two-unit outcome range. This example cannot establish a clinical benefit or the value of ordering a scan.

The two cases are not severity-matched: total motor scores are 31 and 12. Both have unrecorded examination state and recorded LEDD 100 mg/day; identical recorded dose is not equivalent medication exposure. Their MoCA scores (25 and 29) and definitive SAA calls (negative and positive, approximately one month old) are dated biological context, not inputs to these four forecasts. Differences in those markers do not explain the forecast difference causally. The independent matched-pair and multiplicity analyses, including their null results, address different questions and remain in the main results and supplement.

### 5.4 Caudate and putamen reveal regional differences, not a new predictor

The original source rows also contain bilateral caudate binding. Figure 6 displays both regions for all four preselected cases. These values were reconciled to the frozen cohort after selection and do not change the primary putamen model, its prediction, or the definition of a case.

**Figure 6:**
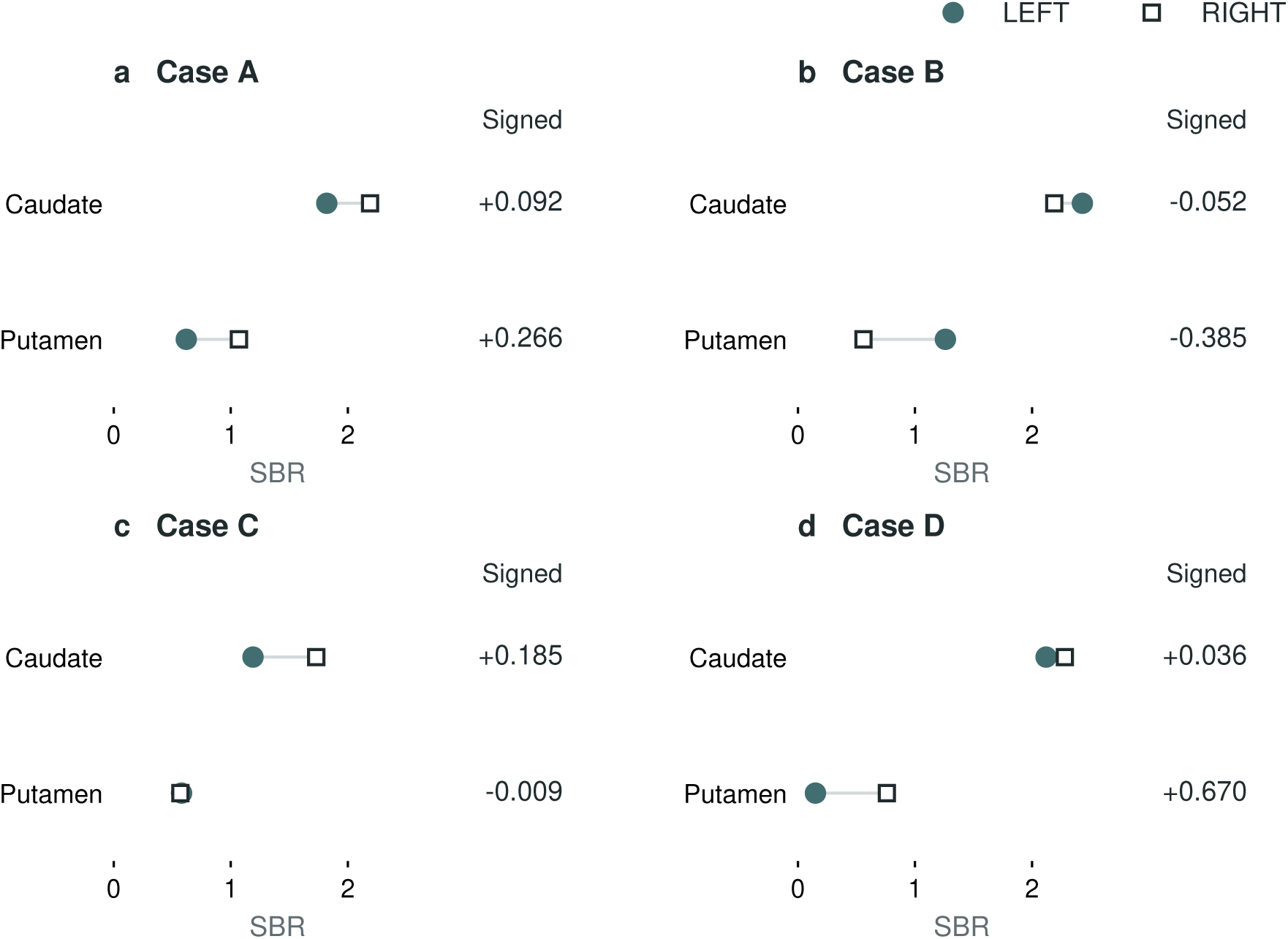
Actual regional evidence behind Cases A–D. Filled circles are left SBR and open squares right SBR; connectors join hemispheres within a region, not measurements over time. The right column gives signed (right-left)/(right+left) ratios. a–b, Putamen and caudate share a sign, but differ in imbalance magnitude. c, Near-symmetric putamen coexists with positive caudate asymmetry. d, Strong positive putamen asymmetry coexists with near-symmetric caudate. All source scans were acquired in the clinical landmark month. Values were read from original SBR analysis rows, not reference brain renders. These are descriptive regional observations, not evidence that post hoc region selection improves prediction.

In Case A, caudate left/right SBRs 1.82/2.19 yield a ratio of 0.0923; in B, 2.43/2.19 yield -0.0519. Both have the same direction as the putamen but less relative imbalance. A reference brain can show the distinct regions; it cannot establish that their measured differences arise from different causal pathways. The scan is a regional tracer-binding summary, not a neuron count, a direct dopamine measurement or a patient-specific tractography reconstruction.

### 5.5 Cases C and D keep weak or conflicting explanations visible

Case C has putamen SBR 0.58/0.57, mean 0.575 and signed ratio -0.0087. Its small asymmetry does not imply preserved overall binding or absence of motor impairment. Left/right motor sums 1/11 give laterality 0.8333 and total motor burden is 17. Caudate SBR 1.19/1.73 gives positive asymmetry 0.1849, but selecting caudate because it looks more concordant in this displayed patient would be a new, post hoc decision. We retain the prespecified putamen model. At follow-up the motor ratio was zero; the signed-model forecast remained approximately 0.538, compared with 0.647 clinically. Neither model resolved the observed loss of side predominance.

Case D supplies the opposite caution. Putamen SBR 0.15/0.76 gives strong positive asymmetry 0.6703, whereas motor sums left/right 12/9 give negative laterality -0.1429. Caudate SBR 2.12/2.28 is nearly symmetric (0.0364). The clinical-only forecast was approximately -0.073 and the signed forecast 0.333; the observed future ratio was 0.143. Preserving sign moved the prediction across zero but overshot. The magnitude-only forecast, approximately 0.006, was closer. A compelling directional story therefore need not identify the best prediction even when it gets the future sign right.

All four cases were chosen from the same strict 32-record intersection without future outcomes or forecast errors. Their selection favored interpretable source availability and prespecified baseline configurations; they are not representative estimates of subgroup prevalence. Keeping the unfavorable and conflicting cases prevents the anatomy from becoming a success-only illustration.

### 5.6 What the longitudinal test adds to an examination

The method does not rediscover that a clinician can see the more affected body side. It asks whether multimodal integration preserves an anatomical coordinate that an undirected representation erases, and whether that coordinate remains associated with future side balance after the current examination is represented. The positive adjusted association addresses the second question; the matched prediction hierarchy estimates the incremental information under specified validation conditions.

The answers are different in strength. The signed anatomical relation reproduces cross-sectionally in S4 using the same orientation. The PPMI adjusted future association survives several robustness tests. The additional prediction gain is small, however, does not improve the binary side decision, and is uncertain under the bounded link and explicit same-state sensitivity. S4 does not validate the longitudinal forecast. These limitations determine the clinical meaning: retain the side, source date, medication documentation, disagreement and uncertainty when constructing a research state, but do not infer global progression rate or treatment choice from its sign.

Taken together, the figures make visible three forms of heterogeneity: opposite but concordant anatomical states; weak directional imaging despite a lateralized examination; and direct imaging– examination conflict. They are distinguishable descriptions of the available evidence, not proven distinct etiologies. A future mechanism-anchored model must explain these patterns and survive external longitudinal testing. The present contribution is the measured coordinate and the tests that keep that ambition separate from the evidence already obtained.

## 6 Data availability

PPMI data were obtained under the PPMI data-use agreement from www.ppmi-info.org/access-data-specimens/download-data (RRID:SCR_006431). S4 data were used under the applicable study data-use terms. Participant-level data and identifiers cannot be redistributed by the authors. Authorized researchers may request the source releases directly from their governing programs. The analysis package available on request provides cohort-construction code, variable definitions, analysis protocols and synthetic tests so that authorized users can reconstruct the derived cohorts locally. Identifier maps and row-level derived results are not redistributed. The outcome-blind illustrative histories display rounded derived measurements under arbitrary one-use labels and relative time; they contain no participant identifier, calendar date or participant scan. No new primary human-participant data were generated.

## 7 Code availability

Analysis code and supporting documentation are maintained at https://github.com/CVC-Lab/pd-signed-laterality. For repository access, please contact the authors. Controlled participant-level data are not redistributed and must be obtained directly from PPMI and S4 under their respective data-use agreements.

## 8 Acknowledgements

This research was supported in part by grants and gifts from the Peter O’Donnell Foundation, the Michael J Fox Foundation, Jim Holland-Backcountry and Michael-Connie Rasor Foundations towards curing Parkinson’s Disease.

Data used in the preparation of this article were obtained on 20 December 2025 from the Parkinson’s Progression Markers Initiative (PPMI) database (www.ppmi-info.org/access-data-specimens/download-data), RRID:SCR_006431. For up-to-date information on the study, visit www.ppmi-info.org. PPMI is a public-private partnership funded by The Michael J. Fox Foundation for Parkinson’s Research and funding partners. The current full funding-partner list for this release includes 4D Pharma, AbbVie, AcureX, Allergan, Amathus Therapeutics, Aligning Science Across Parkinson’s, AskBio, Avid Radiopharmaceuticals, BIAL, BioArctic, Biogen, Biohaven, BioLegend, BlueRock Therapeutics, Bristol Myers Squibb, Calico Labs, Capsida Biotherapeutics, Celgene, Cerevel Therapeutics, Coave Therapeutics, DaCapo Brainscience, Denali, Edmond J. Safra Foundation, Eli Lilly, Gain Therapeutics, GE HealthCare, Genentech, GSK, Golub Capital, Handl Therapeutics, insitro, Jazz Pharmaceuticals, Johnson & Johnson Innovative Medicine, Lundbeck, Merck, Meso Scale Discovery, Mission Therapeutics, Neurocrine Biosciences, Neuron23, Neuropore, Pfizer, Piramal, Prevail Therapeutics, Roche, Sanofi, Servier, Sun Pharma Advanced Research Company, Takeda, Teva, UCB, Vanqua Bio, Verily, Voyager Therapeutics, the Weston Family Foundation and Yumanity Therapeutics.

We thank the participants and investigators of PPMI and S4. Computational analyses used resources of the Texas Advanced Computing Center at The University of Texas at Austin. The PPMI and S4 investigators did not participate in the present analysis or approve the manuscript’s conclusions.

## 9 Author contributions

H.T. and P.Y. contributed equally. H.T.: conceptualization, methodology, software, formal analysis, validation, visualization, writing: original draft, and writing: review and editing. P.Y.: conceptualization, data curation, methodology, software, formal analysis, validation, visualization, writing: original draft, and writing: review and editing. C.B.: conceptualization, methodology, supervision, project administration, resources, and writing: review and editing. All authors reviewed and approved the manuscript.

## Supplementary Information

This Supplementary Information reports the complete secondary results, patient-specific figures and sensitivity analyses for the signed-laterality study. Source provenance, variable definitions, sign conventions, timing rules and the prespecified primary analysis are reported in the main manuscript.

### Supplementary Results

#### S1. Longitudinal cohort and participant-held-out performance

| Flow element | Participants |
| --- | --- |
| PPMI participants classified as PD with dated motor data | 1,602 |
| Valid analyzed left-right putamen SBR | 1,439 |
| Valid scan-clinical landmark and 12-month outcome | 1,204 |
| 24-month outcome available | 901 |
| 36-month outcome available | 682 |

| Scan-clinical separation | Participants | Percent of primary cohort |
| --- | --- | --- |
| Same calendar month | 636 | 52.8% |
| 1 month | 284 | 23.6% |
| 2 months | 204 | 16.9% |
| 3 months | 80 | 6.6% |

| Baseline examination state | Participants | Percent |
| --- | --- | --- |
| Explicit OFF | 131 | 10.9% |
| Explicit ON | 51 | 4.2% |
| Unknown or ambiguous | 1,022 | 84.9% |

The one-landmark design removes ambiguity between participant count and repeated prediction opportunities. The large unknown-state cell motivated mandatory reporting of the explicit same-state sensitivity rather than treating state adjustment as complete.

| Horizon | Model | n | $R^2$ | MAE | RMSE | Spearman r | Side accuracy |
| --- | --- | --- | --- | --- | --- | --- | --- |
| 12 | Clinical state | 1,204 | 0.713 | 0.234 | 0.309 | 0.853 | 0.902 |
| 12 | Overall DaT signal | 1,204 | 0.712 | 0.234 | 0.309 | 0.853 | 0.899 |
| 12 | Magnitude-only imaging | 1,204 | 0.712 | 0.233 | 0.309 | 0.853 | 0.902 |
| 12 | Direction-aware imaging | 1,204 | 0.730 | 0.227 | 0.300 | 0.863 | 0.898 |
| 24 | Clinical state | 901 | 0.675 | 0.242 | 0.318 | 0.835 | 0.914 |
| 24 | Overall DaT signal | 901 | 0.673 | 0.242 | 0.319 | 0.834 | 0.912 |

| Horizon | Model | n | $R^2$ | MAE | RMSE | Spearman<br>r | Side<br>accuracy |
| --- | --- | --- | --- | --- | --- | --- | --- |
| 24 | Magnitude-only imaging | 901 | 0.675 | 0.241 | 0.318 | 0.834 | 0.912 |
| 24 | Direction-aware imaging | 901 | 0.681 | 0.238 | 0.315 | 0.840 | 0.916 |
| 36 | Clinical state | 682 | 0.671 | 0.225 | 0.290 | 0.835 | 0.887 |
| 36 | Overall DaT signal | 682 | 0.671 | 0.225 | 0.289 | 0.836 | 0.887 |
| 36 | Magnitude-only imaging | 682 | 0.672 | 0.225 | 0.289 | 0.837 | 0.887 |
| 36 | Direction-aware imaging | 682 | 0.676 | 0.223 | 0.288 | 0.840 | 0.889 |

The future-side classification column is descriptive. A tied future laterality has no affected side and was omitted from accuracy. No classification threshold was tuned.

| Horizon | Reference | $\Delta R^2$ (95% CI) | Delta MAE (95% CI) | Delta side accuracy (95% CI) |
| --- | --- | --- | --- | --- |
| 12 | Clinical state | 0.016 (0.007, 0.026) | -0.0064 (-0.0109, -0.0021) | -0.003 (-0.014, 0.007) |
| 12 | Overall DaT signal | 0.018 (0.008, 0.027) | -0.0067 (-0.0111, -0.0023) | -0.001 (-0.011, 0.009) |
| 12 | Magnitude-only imaging | 0.018 (0.009, 0.028) | -0.0063 (-0.0107, -0.0019) | -0.003 (-0.013, 0.006) |
| 24 | Overall DaT signal | 0.008 (-0.001, 0.018) | -0.0041 (-0.0083, 0.0001) | 0.003 (-0.007, 0.014) |
| 36 | Overall DaT signal | 0.004 (-0.005, 0.013) | -0.0019 (-0.0056, 0.0020) | 0.002 (-0.009, 0.012) |

All intervals use 10,000 paired participant bootstraps. Only the 12-month contrasts supported a linear increment; later-horizon intervals did not exclude no increment.

The fixed-prediction intervals above condition on one set of out-of-fold fits. A separate model-refitting sensitivity, specified after the primary analysis, therefore refitted the complete imputation, scaling and ridge pipeline in 1,000 participant bootstrap samples and evaluated each fit only on participants absent from training.

| Reference | Median $\Delta R^2$ (2.5th, 97.5th percentiles) | Median delta MAE (2.5th, 97.5th percentiles) | $\Pr(\Delta R^2 > 0)$ | $\Pr(\text{delta MAE} < 0)$ |
| --- | --- | --- | --- | --- |
| Clinical state | 0.0159 (-0.0002, 0.0276) | -0.0061 (-0.0124, 0.0011) | 0.973 | 0.958 |
| Overall DaT signal | 0.0175 (0.0015, 0.0292) | -0.0068 (-0.0129, 0.0000) | 0.981 | 0.973 |
| Magnitude-only imaging | 0.0171 (0.0013, 0.0295) | -0.0063 (-0.0127, 0.0007) | 0.982 | 0.958 |

These percentile ranges include resampled participant composition and complete pipeline refitting.

They support split stability but are not independent-cohort validation intervals.

#### S2. Robustness, calibration and independent replication

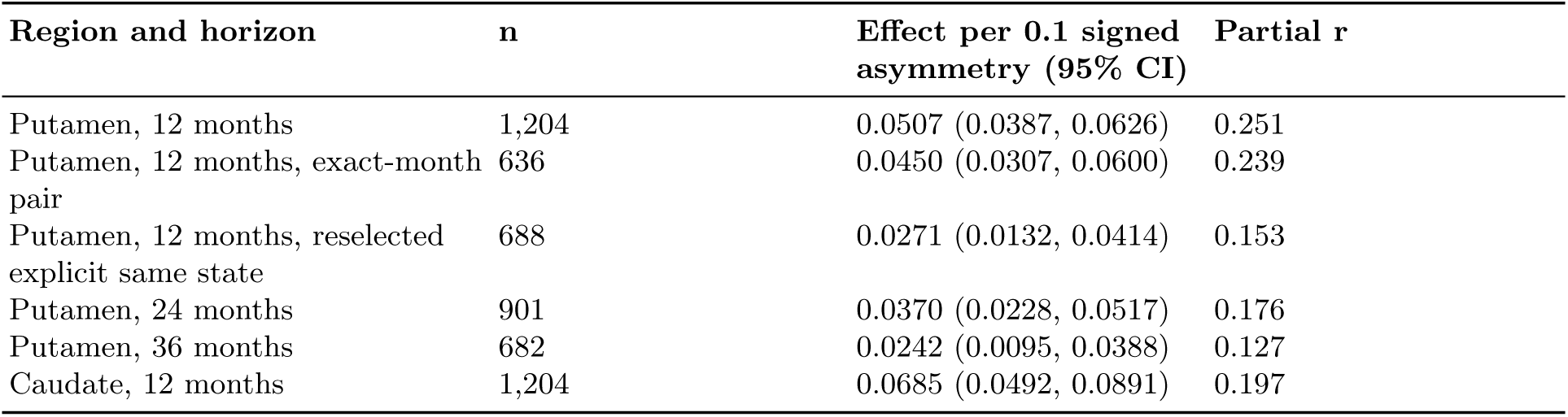

The unnormalized 12-month sensitivity estimated 0.757 additional right-minus-left paired motor points per 0.1 signed putamen asymmetry (95% CI 0.600-0.918). This agrees in sign with the normalized endpoint and shows that the association is not created solely by the ratio. The non-lateralized gait-plus-freezing negative-control coefficient was -0.038 per unit signed putamen asymmetry (95% CI -0.328 to 0.251; n=1,204).

| Evaluation and reference | Test or OOF n | $\Delta R^2$ (95% CI) | Delta MAE (95% CI) |
| --- | --- | --- | --- |
| Site-disjoint signed versus overall DaT | 977 | 0.0146 (0.0056, 0.0238) | -0.0062 (-0.0107, -0.0017) |
| Strict calendar signed versus overall DaT | 582 | 0.0191 (0.0066, 0.0325) | -0.0076 (-0.0135, -0.0019) |
| Strict calendar signed versus clinical state | 582 | 0.0087 (-0.0066, 0.0248) | -0.0022 (-0.0095, 0.0051) |
| Explicit same-state signed versus overall DaT | 688 | 0.0063 (-0.0023, 0.0148) | -0.0015 (-0.0053, 0.0022) |

The strict-calendar advantage over clinical state and every explicit same-state prediction interval included no increment. These tests support a directional association and a modest contrast with non-directional imaging; they do not establish deployment-ready prediction.

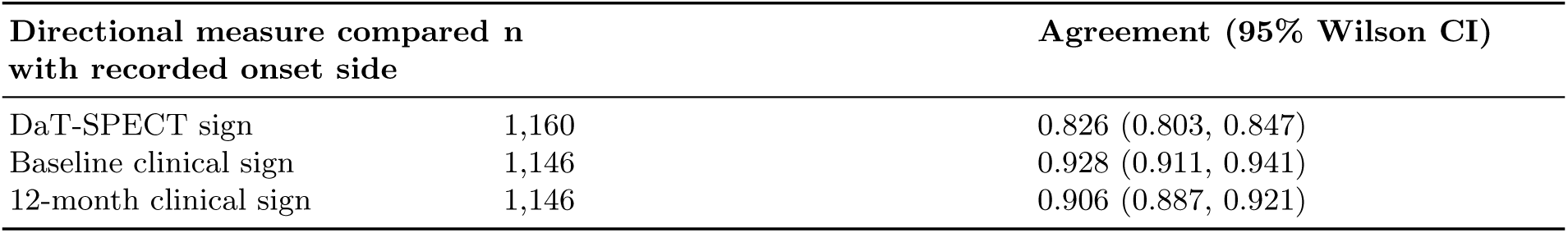

Adding lateralized onset side to the prespecified adjustment set retained an imaging effect of 0.0397 per 0.1 signed asymmetry (95% CI 0.0282-0.0518; n=1,169). Adding recorded handedness yielded 0.0398 (0.0281-0.0519; n=1,168). These are biological triangulation checks, not independent validation.

| Maximum DaT-clinical gap | Participants | Signed Pearson r (95% CI) | Signed Spearman rho | Magnitude Pearson r (95% CI) |
| --- | --- | --- | --- | --- |
| Exact month | 1,059 | 0.665 (0.632, 0.696) | 0.681 | 0.202 (0.143, 0.263) |
| 1 month | 1,223 | 0.674 (0.645, 0.702) | 0.692 | 0.192 (0.136, 0.247) |
| 3 months | 1,367 | 0.676 (0.649, 0.702) | 0.692 | 0.206 (0.153, 0.258) |

| S4 analysis | n | Pearson r (95% CI) | Spearman rho | Leave-one-out r range | Winsorized r |
| --- | --- | --- | --- | --- | --- |
| Putamen, OFF-state primary | 56 | 0.551 (0.363, 0.701) | 0.590 | 0.525 to 0.614 | 0.554 |
| Caudate, OFF-state secondary | 56 | 0.664 (0.562, 0.753) | 0.654 | 0.651 to 0.682 | 0.671 |
| Putamen, ON-state sensitivity | 38 | 0.401 (0.159, 0.609) | 0.469 | 0.339 to 0.457 | 0.397 |
| Absolute putamen versus absolute OFF motor | 56 | 0.058 (-0.182, 0.294) | 0.046 | 0.006 to 0.120 | 0.051 |

The median absolute S4 scan-to-UPDRS separation was 17 days (IQR 8-42). The primary PPMI and S4 coefficients were not detectably different by Fisher’s transformation (z=1.449, p=0.147). This test assesses incompatibility, not cohort equivalence.

| Evaluation | Model | n | Calibration intercept | Calibration slope | Median AE |
| --- | --- | --- | --- | --- | --- |
| Participant-held-out | Clinical state | 1,204 | -0.001 | 1.026 | 0.191 |
| Participant-held-out | Direction-aware | 1,204 | -0.002 | 1.016 | 0.185 |
| Strict calendar | Clinical state | 582 | -0.006 | 1.119 | 0.183 |
| Strict calendar | Direction-aware | 582 | -0.062 | 1.083 | 0.181 |
| Explicit same state | Clinical state | 688 | 0.001 | 1.032 | 0.163 |
| Explicit same state | Direction-aware | 688 | 0.001 | 1.026 | 0.164 |

The strict-calendar direction-aware model overpredicted the mean signed outcome (predicted 0.072 versus observed 0.015), despite retaining an increment over non-directional imaging. No recalibration was applied.

**Supplementary Figure 1:**
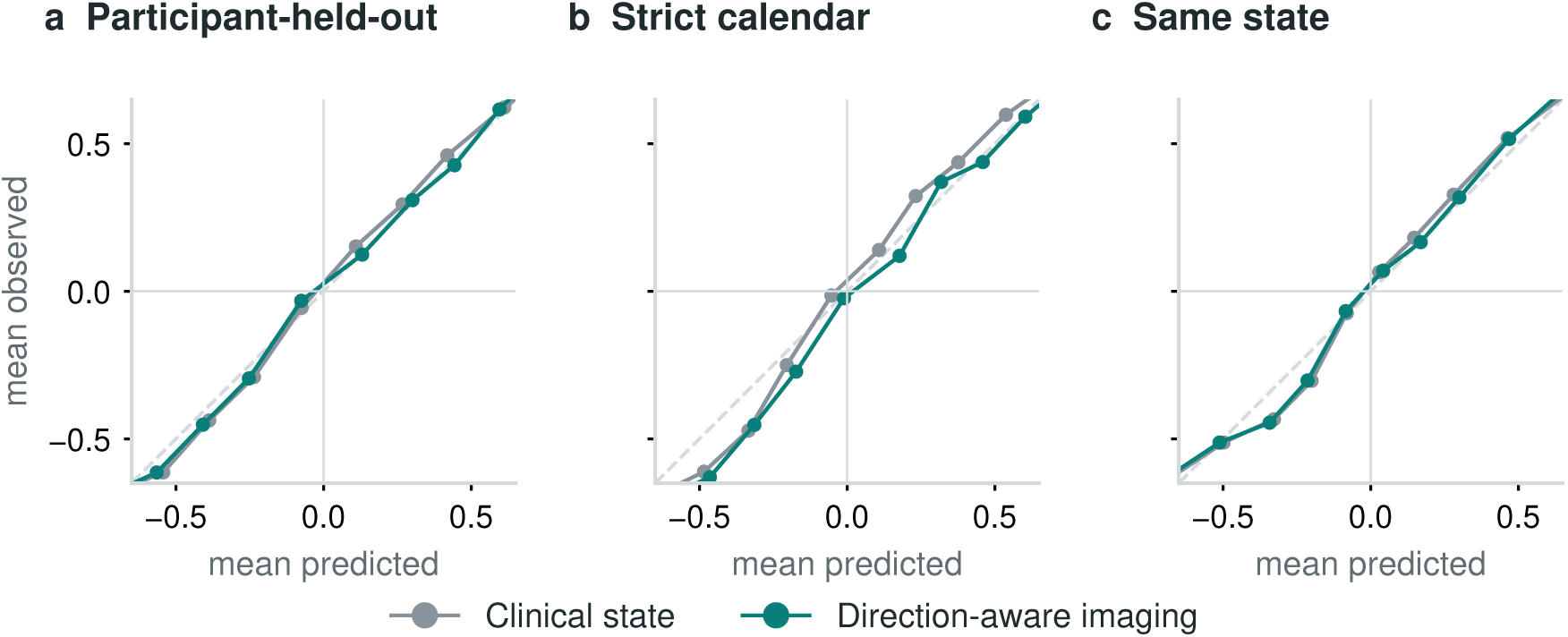
Calibration of future signed laterality. Decile means for a, participant-held-out (n=1,204), b, strict-calendar (n=582 test participants) and c, retrospective explicit same-state (n=688) predictions. Within each evaluation, participants were divided into ten groups by predicted value; points show the mean predicted and observed signed motor laterality in each group, and connecting lines aid reading. Gray is the clinical-state model and teal is the direction-aware model. The dashed diagonal denotes ideal calibration, while horizontal and vertical zero lines separate left- and right-dominant values. These are descriptive summaries of held-out or untouched predictions; no calibration model was fitted and no recalibration was applied.

#### S3. Outcome-blind patient-specific visualizations

The outcome-blind patient-history protocol was specified before any illustrative participant was selected or viewed. Eligibility required a same-month DaT-SPECT and clinical landmark; MoCA, definitive SAA, CSF alpha-synuclein and NfL observed no later than the landmark and no more than 12 months old; treatment context; and all four participant-held-out forecasts. The strict intersection contained 32 participants.

Across the full 1,204-participant cohort, nominal 90% conformalized quantile intervals achieved 95.5% empirical coverage and had median width 1.504 on the bounded -1 to 1 outcome scale. Only one of the two concordant directional examples retained the imaging-indicated sign at 12 months. The records therefore demonstrate current-state resolution and uncertainty, not four retrospectively curated forecast successes.

The main patient-history figure uses arbitrary one-use Case A–B labels, and the four supplementary dossiers use one-use Case A–D labels, relative months and rounded values. They contain no participant identifier or calendar date. The internal mapping and row-level derived tables remain controlled; case-level and aggregate figures are governed by the same program data-use and publication requirements.

Each dossier separates dated source availability, bilateral state, observed history and held-out prediction. Filled blue and open rust marks encode left and right, respectively; they are not severity colors. Clinical measurements, demographics, examination state and LEDD enter the clinical model; MoCA, SAA and CSF are display context only. The forecast panel compares all four ridge models and separately shows the quantile model’s median with its own conformal interval. No interval is transferred between models. Population-level inference comes from the complete cohorts and bootstrap analyses, not from these four records alone.

**Supplementary Figure 2:**
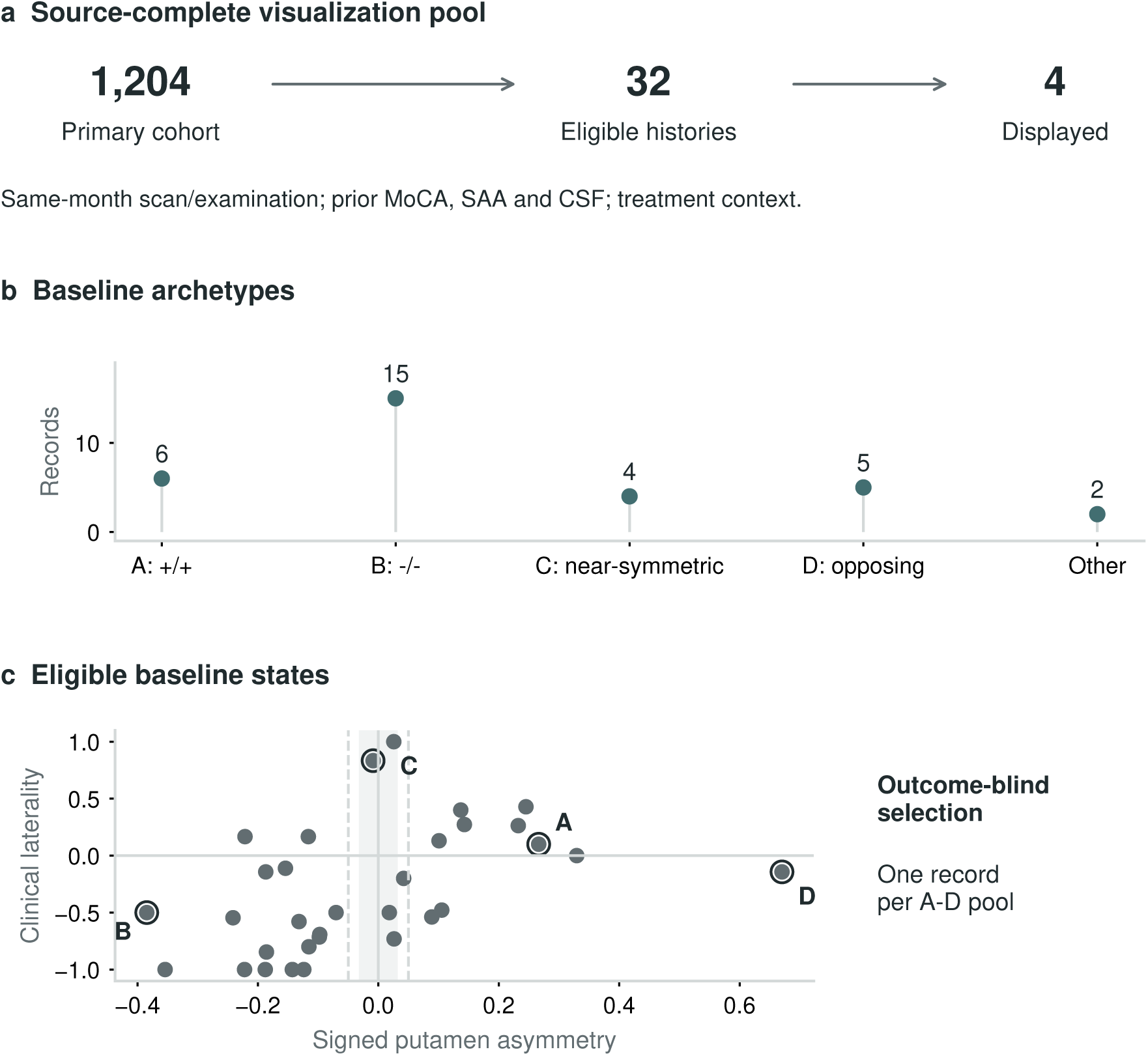
Outcome-blind selection of actual multimodal histories. a, A prespecified source-completeness criterion narrowed the 1,204-participant future-only cohort to 32 strict visualization candidates (2.7%). b, Mutually exclusive baseline definitions yielded six positive-sign concordant records (A), 15 negative-sign mirror records (B), four near-symmetric imaging records (C), five records with opposing imaging and clinical signs (D), and two records outside these archetypes. One record from each A–D pool was ranked using landmark evidence, interval width, model disagreement and source freshness; remaining ties were resolved deterministically without outcome information. c, All 32 candidates are shown in the signed DaT–motor state plane; outlined letters identify the four displayed records. The shaded band marks absolute DaT asymmetry no greater than 0.03 and dashed lines mark magnitudes of 0.05. Future value, sign, error and trajectory were excluded. Counts describe the visualization pool, not population prevalence.

**Supplementary Figure 3:**
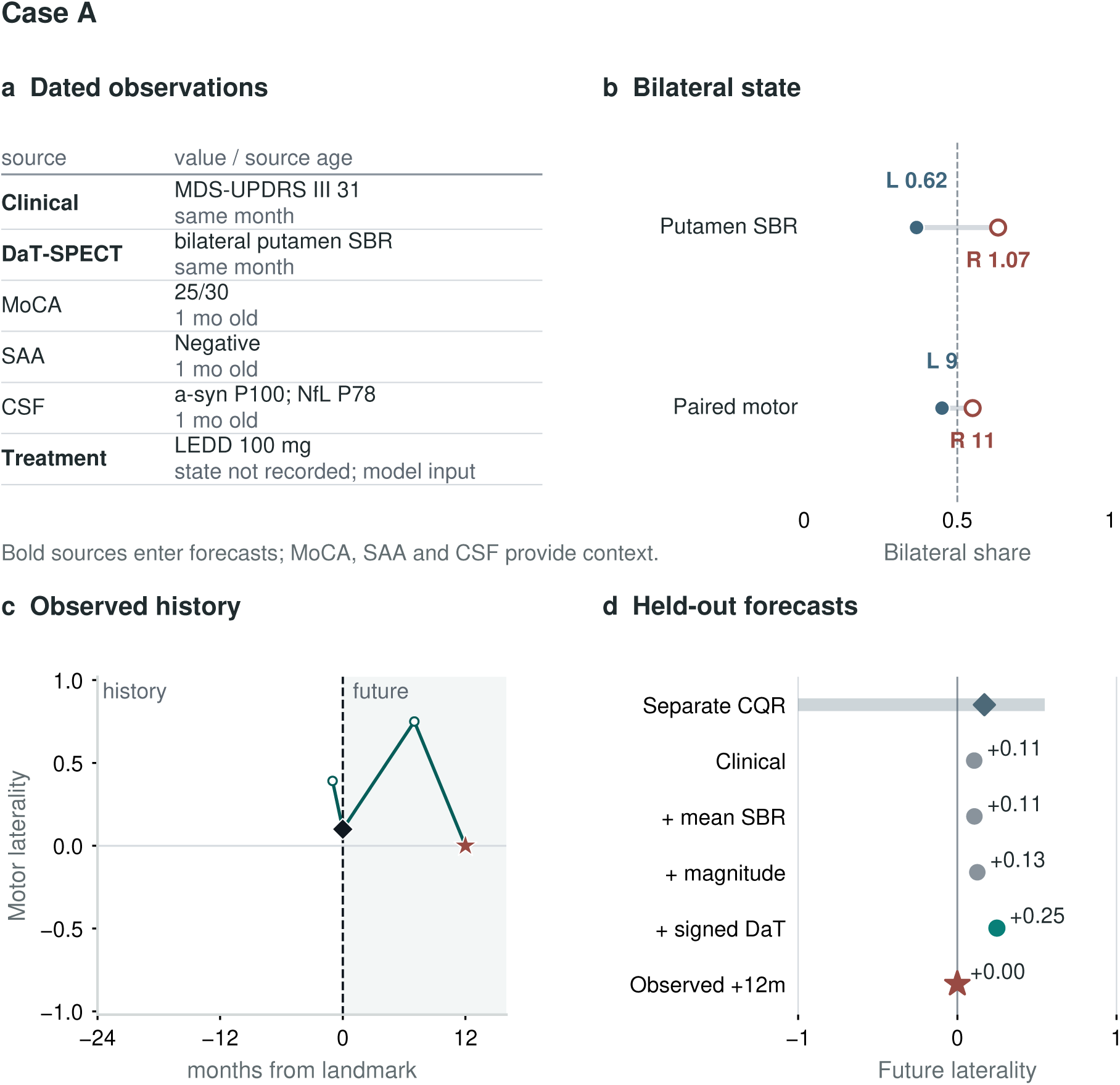
Case A: a concordant current state did not ensure persistent future direction. One participant from the 32-record pool had lower left putamen binding and greater right-body motor burden at the landmark. a, dated sources and their model roles; CSF percentiles refer to the visualization pool, not clinical reference ranges. b, filled blue and open rust marks denote left and right bilateral shares; the dashed line is equal share. c, circles and lines show measured laterality at actual relative months, the diamond is the landmark and the star the selected future observation. d, gray clinical, overall-SBR and magnitude-only predictions are compared with the teal signed-SBR prediction and the observed star. The separate conformalized quantile regression (CQR) diamond and shaded 90% interval belong to the quantile model alone. Signed imaging increased the forecast to 0.249, whereas the observed laterality was zero. Selection excluded this future outcome and error; the record is not evidence of individual benefit.

**Supplementary Figure 4:**
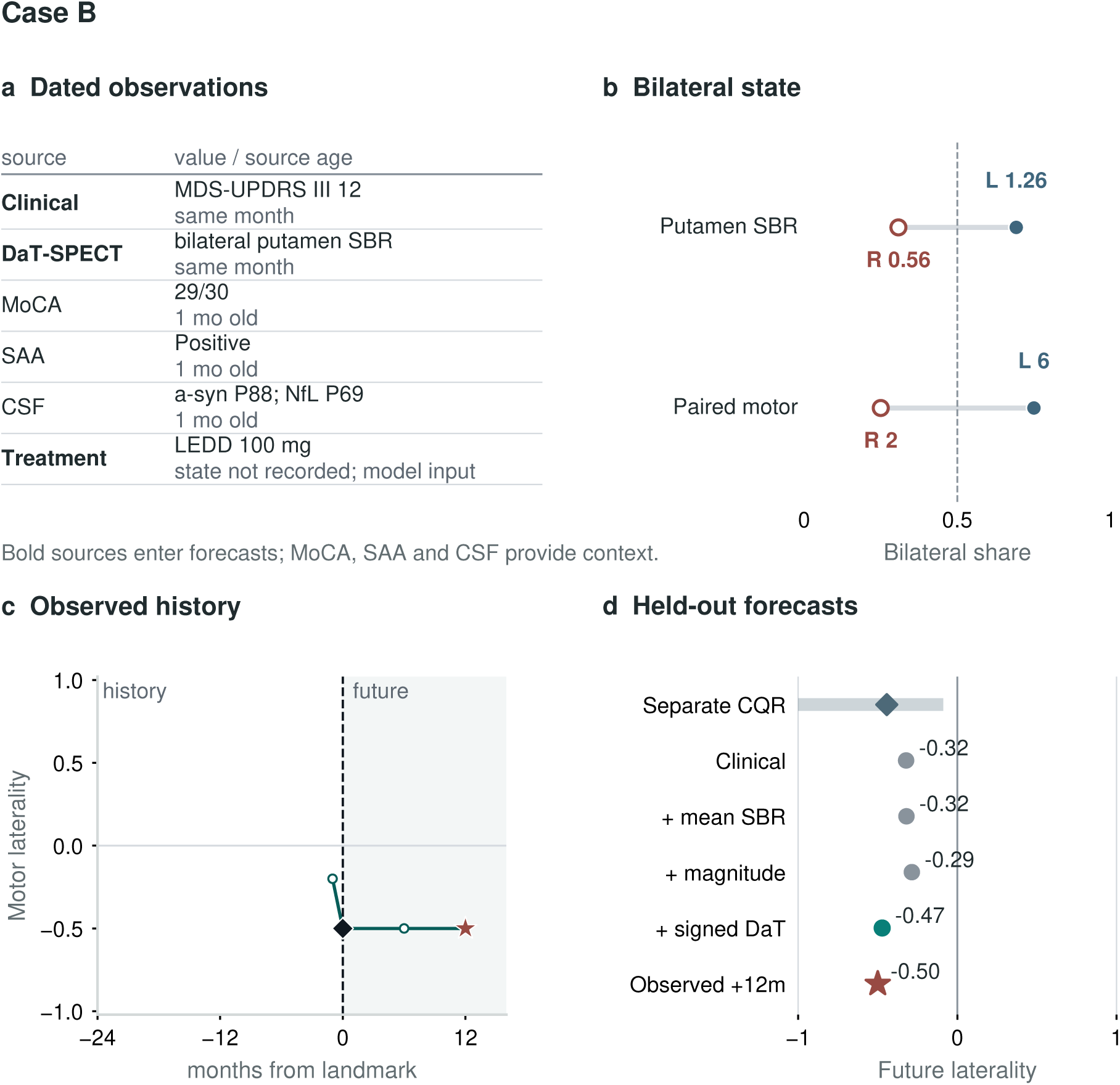
Case B: concordant left-body predominance persisted. One participant from the 32-record pool had lower right putamen binding and greater left-body motor burden. a, dated sources and model roles; CSF percentiles are descriptive within-pool ranks. b, filled blue and open rust marks denote left and right shares, with equality dashed. c, measured history at actual relative months, the landmark diamond and the future observation star. d, comparison of all four ridge forecasts with the observed laterality of -0.500. Signed imaging yielded -0.472, closer than the clinical forecast of -0.322. The separate conformalized quantile regression (CQR) diamond and shaded 90% interval belong to the quantile model, not the ridge predictions. The record was selected without its future outcome or error; this favorable example does not establish patient-level treatment utility.

**Supplementary Figure 5:**
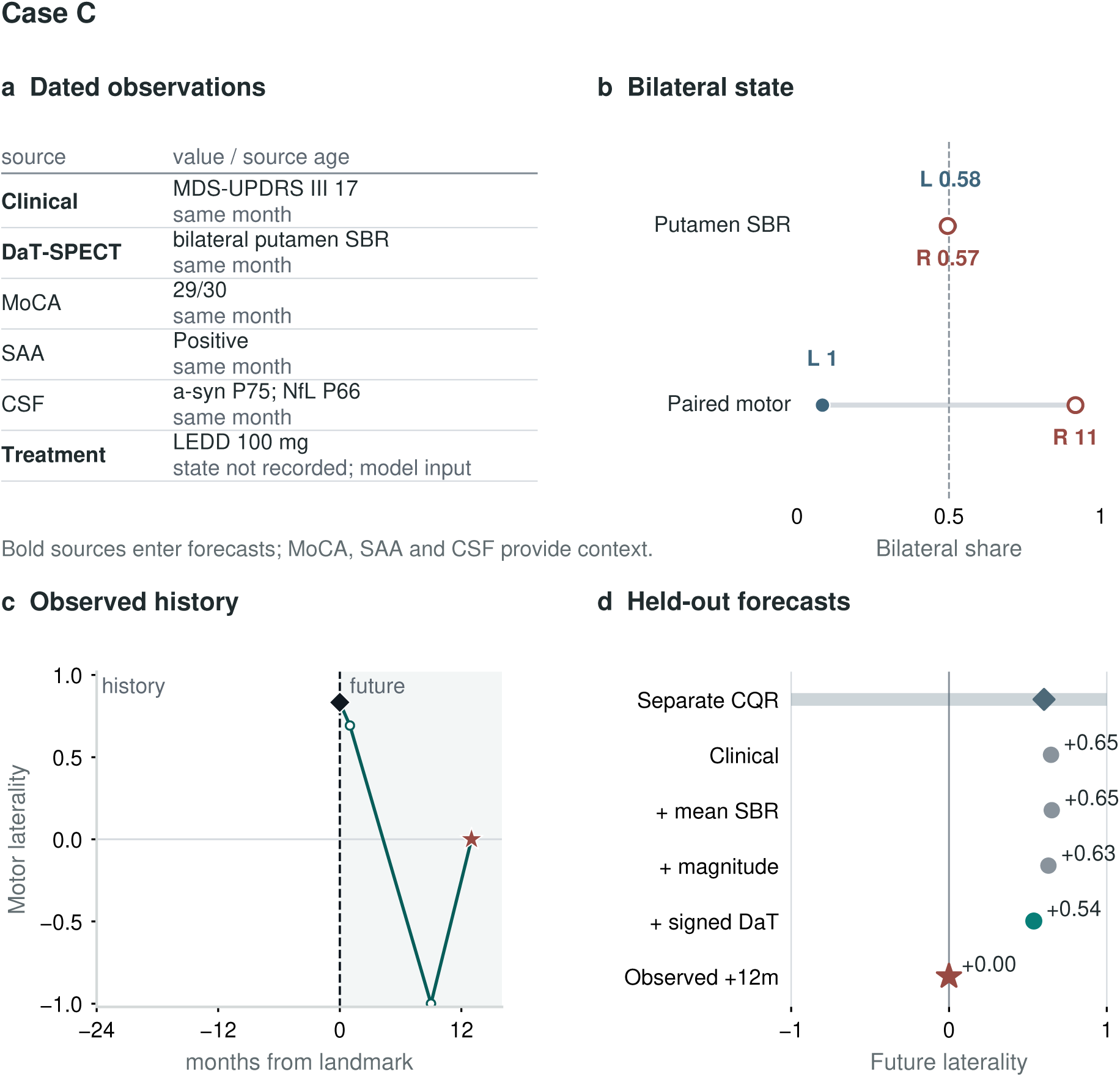
Case C: little imaging direction despite marked clinical laterality. One participant from the 32-record pool had almost equal putamen SBR but predominantly right-body motor signs. a, dated sources and model roles; b, filled blue left and open rust right shares against dashed equality. c, only actual observations, with the landmark diamond and selected future star. d, comparison of the four ridge forecasts and, separately, the quantile-model median and its shaded 90% conformal interval. Signed imaging reduced the clinical forecast from 0.647 to 0.538, but the observed laterality was zero. The broad interval and remaining error preserve the unresolved future state; near-symmetry is not a distinct molecular diagnosis. Selection excluded the participant’s future outcome and error.

**Supplementary Figure 6:**
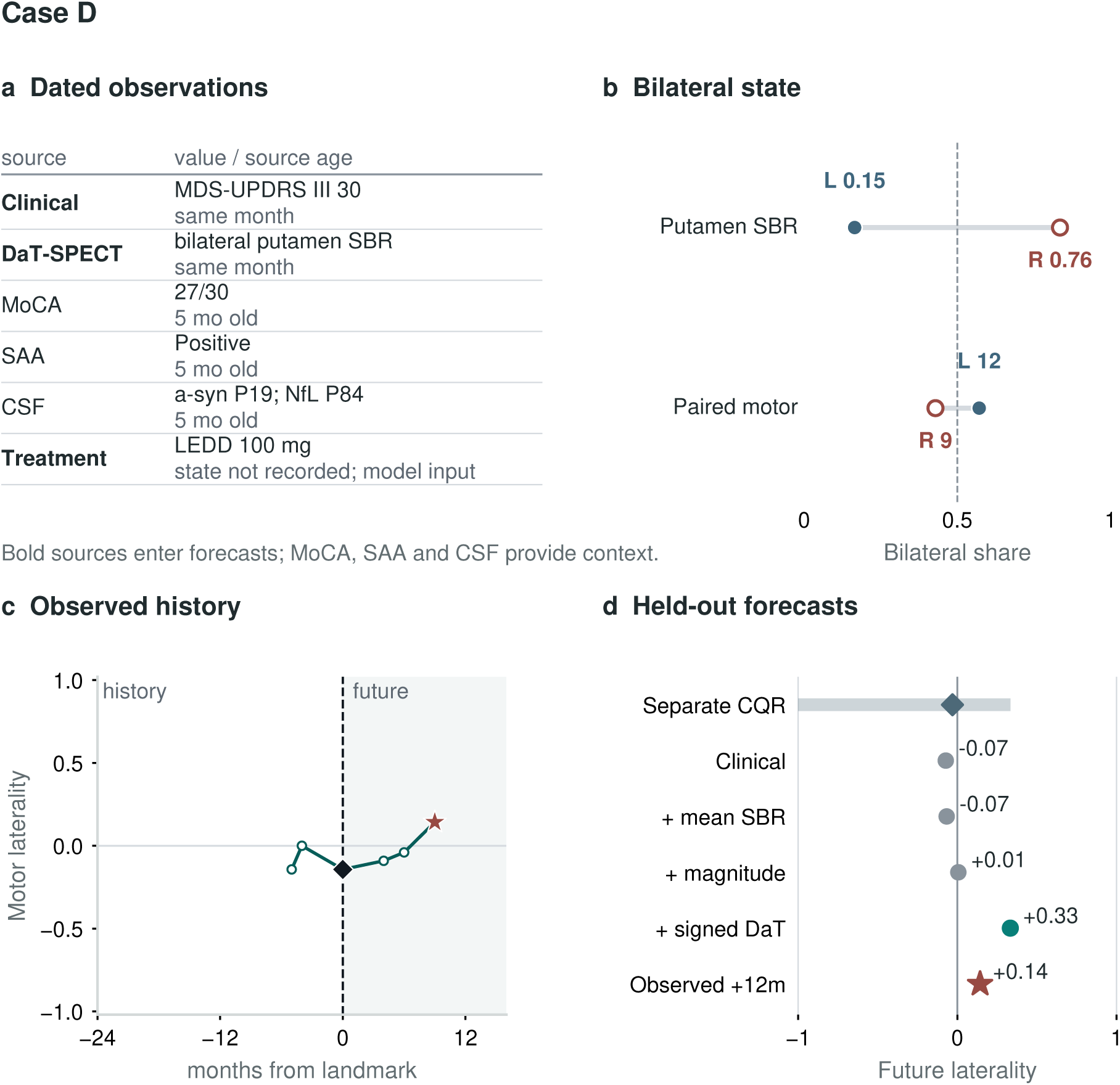
Case D: imaging–clinical conflict must remain visible. One participant from the 32-record pool had lower left putamen binding but greater left-body motor burden. a, dated sources and model roles; b, filled blue left and open rust right shares against dashed equality. c, actual observed history, the landmark diamond and selected future star. d, the clinical, overall-SBR, magnitude-only and signed-SBR ridge forecasts are compared with the observed 0.143 laterality. Signed imaging changed the prediction from -0.073 to 0.333, matching the future sign but overshooting its magnitude; magnitude-only imaging was closer to the observed value. The conformalized quantile regression (CQR) median and shaded 90% interval are a separate model’s output. Selection excluded the future outcome and error. The figure documents conflict and uncertainty, not causal resolution.

**Supplementary Figure 7:**
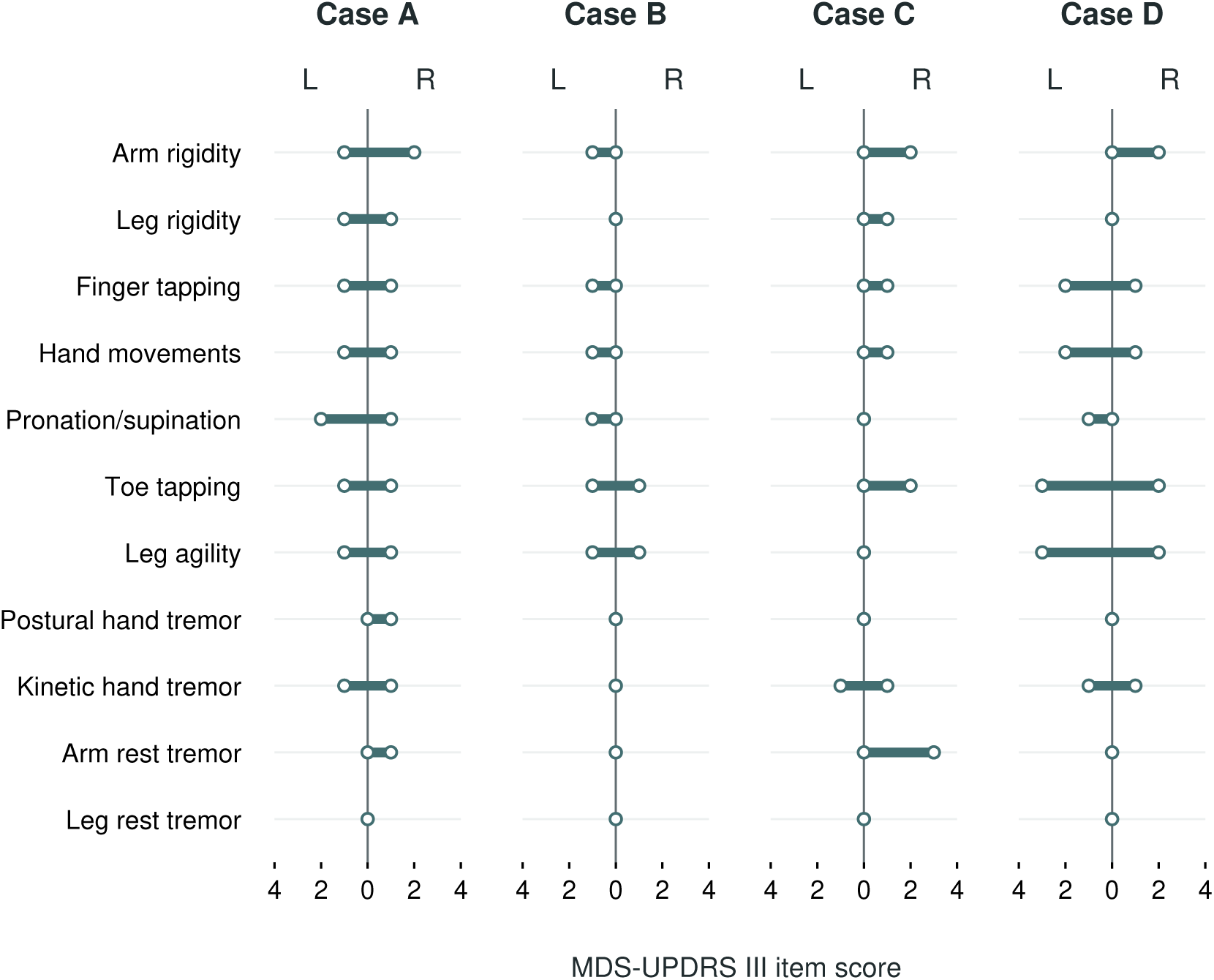
Complete paired examination profiles behind Cases A–D. Each row shows one observed MDS-UPDRS Part III domain on a mirrored 0–4 scale: the left-side score extends left of zero and the right-side score extends right. Endpoints are observed ordinal scores, not signed severity, inferred symptoms or reconstructed motion; a point at zero denotes a recorded zero. Each selected landmark contained one source examination, and the item values reproduce the frozen side totals (A: L/R 9/11; B: 6/2; C: 1/11; D: 12/9). A/B have opposite concordant imaging directions; C has near-symmetric imaging and D imaging–clinical disagreement. Keeping all 11 domains visible prevents a greater aggregate side score from being mistaken for uniform worsening of every symptom on that side. The examination state was unrecorded in all four examples.

#### S4. Directional-disagreement robustness

The prespecified discordance analysis used one landmark per participant and a continuous primary exposure. Categorical patterns were retained for clinical readability rather than used to choose the inferential result. Every bootstrap draw resampled participants and refitted imputation, scaling, site encoding and the complete adjusted model.

| Analysis | n | Stable side | Adjusted odds ratio (95% CI) |
| --- | --- | --- | --- |
| Continuous disagreement, primary | 1,178 | 89.4% | 0.491 (0.297, 0.711) per IQR-standardized unit |
| Continuous disagreement, future ties excluded | 1,153 | 91.3% | 0.462 (0.257, 0.703) per IQR-standardized unit |
| Continuous disagreement, exact-month pairing | 624 | 89.3% | 0.425 (0.191, 0.689) per IQR-standardized unit |
| Continuous disagreement, explicit same medication state | 138 | 85.5% | 1.148 (numerically unstable CI) per IQR-standardized unit |
| Discordant versus concordant, threshold 0.02 | 1,098 | 90.3% | 0.412 (0.198, 0.705) |
| Discordant versus concordant, threshold 0.03 | 1,056 | 90.4% | 0.335 (0.152, 0.581) |
| Discordant versus concordant, threshold 0.05 | 988 | 91.1% | 0.242 (0.092, 0.428) |

The explicit same-state subset is non-confirmatory: only 138 records remained and sparse separation produced a 95% bootstrap CI from 2.1 × 10*^−^*^7^ to 2.05 × 10^9^. Continuous disagreement also increased absolute out-of-fold prediction error by 0.0212 laterality units per IQR-standardized unit (95% CI 0.0057 to 0.0377). A discordance-informed ranker changed AURC only from 0.179 to 0.178 relative to clinical uncertainty, so no selective-prediction claim was supported.

After requiring absolute baseline clinical laterality above 0, 0.05 and 0.10, the adjusted continuous-disagreement odds ratios were 0.433 (95% CI 0.239-0.667; n=1,038), 0.428 (0.223-0.683; n=1,008) and 0.417 (0.186-0.699; n=927), respectively. A participant-clustered repeated-horizon model containing 2,554 observations from 1,147 participants at 12, 24 and 36 months yielded OR=0.629 per IQR-standardized unit (0.425-0.895). These are robustness analyses of one association, not independent replications.

#### S5. Medication-state selection and bounded-outcome sensitivity

The explicit same-state analysis was not a random subsample. It selected participants with observed future examination state and substantially different duration and treatment-record availability. Standardized mean differences (SMDs) compare the explicit same-state cohort with the primarycohort [utbl1]

| Characteristic | Primary cohort | Explicit same state | Primary unknown state | SMD: explicit minus unknown |
| --- | --- | --- | --- | --- |
| Age, years | 63.06 | 63.79 | 62.98 | 0.083 |
| Disease duration, years | 1.26 | 2.70 | 0.91 | 1.173 |
| Baseline MDS-UPDRS III | 22.14 | 25.48 | 21.68 | 0.347 |
| Signed putamen asymmetry | 0.018 | 0.006 | 0.020 | -0.071 |
| Baseline clinical laterality | 0.039 | 0.001 | 0.059 | -0.095 |
| LEDD history observed, % | 26.0 | 99.6 | 13.0 | 3.568 |

The primary cohort contained 131 OFF, 51 ON and 1,022 unknown baseline states. The reselected cohort contained 532 OFF–OFF and 156 ON–ON records. Its persistence of the adjusted signed association is informative, but the strong duration and LEDD-selection differences prevent population-representative or medication-invariant interpretation.

| Outcome model | Participants | Signed versus overall<br>$\Delta R^2$ (95% CI) | Delta MAE (95% CI) |
| --- | --- | --- | --- |
| Primary linear | 1,204 | 0.0176 (0.0085, 0.0267) | -0.0067 (-0.0111, -0.0022) |
| Primary linear clipped to [-1,1] | 1,204 | 0.0176 (0.0087, 0.0269) | -0.0067 (-0.0109, -0.0025) |
| Logit-transformed bounded<br>ridge | 1,204 | 0.0041 (-0.0057, 0.0140) | 0.0038 (-0.0007, 0.0081) |

No primary held-out linear prediction exceeded [-1,1], so clipping changed no value. In the separate adjusted fractional-logit association model, the odds ratio per 0.1 signed putamen asymmetry was 1.126 (robust 95% CI 1.091–1.161). Thus the association survives a bounded mean model, whereas the small incremental prediction depends on the outcome link and is not supported as robust prognostic utility. No post-validation recalibration was applied.

#### S6. Complete dated molecular and cognitive multiplicity family

Context was restricted to measurements observed at or before the landmark and linked within the preceding 12 months, without filling unmeasured assays. Eleven comparisons formed one prespecified Benjamini–Hochberg family. For continuous measures, effect is the Mann–Whitney probability-of-superiority minus 0.5; for SAA it is the Fisher odds ratio. None passed q<0.05.

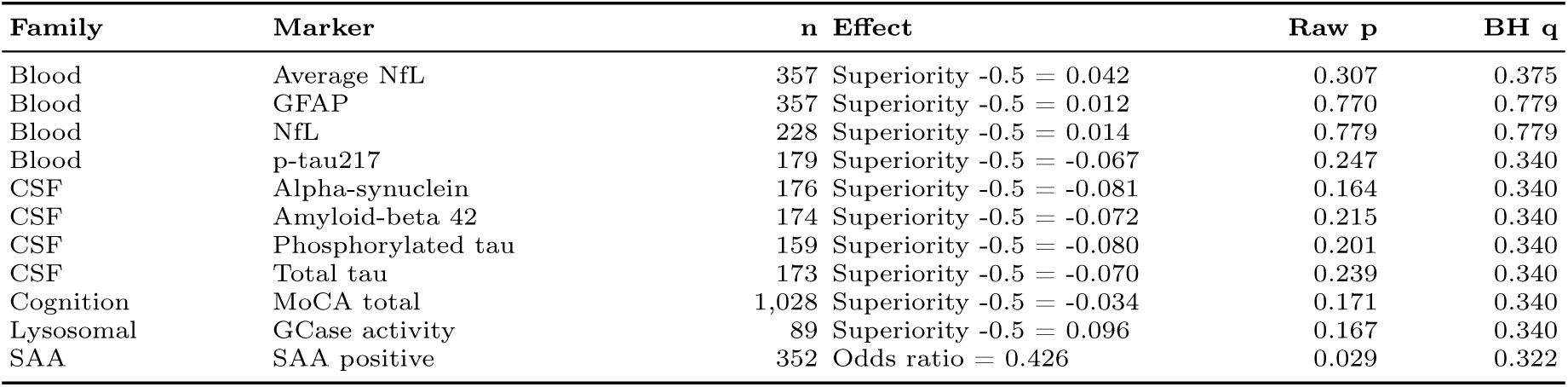

SAA positivity was 77.2% among 57 discordant records and 88.8% among 295 concordant records. Its unadjusted p value was the smallest in the family, but q=0.322. This complete table prevents selective emphasis on that nominal result. The screen supplies secondary context only and cannot define a molecular subtype.

#### S7. Similar summaries, directional information and the limits of molecular explanation

The September descriptive sign-loss audit used the primary 1,204-person cohort without refitting any predictive model. Of 1,056 eligible records with non-zero clinical laterality and absolute putamen asymmetry above 0.03, 2,959 candidate edges met the fixed motor-total, mean-SBR and magnitude calipers. Deterministic greedy matching retained 365 non-overlapping pairs (730 participants) with opposite imaging signs. Clinical signs were also opposite in 281 pairs. This is conditional on the stated matching rules, not an estimate of population prevalence or a diagnosis error rate. The clinical reference already retains clinical direction.

A different, earlier analysis asked whether secondary biological-profile differences explained future divergence among clinically and imaging-similar participants, using the matching and resampling procedures in main Methods. The complete profile comprised MoCA, SAA, CSF alpha-synuclein and NfL. Neither that complete profile nor the reduced-coverage comparisons showed a clear relation to future laterality divergence. All intervals below use the completed 50,000-resample analyses.

Table 17: Median absolute difference between paired held-out forecasts in the 365 descriptive pairs. Separation measures how predictions differ; it is not forecast accuracy.

| Model | Median within-pair difference |
| --- | --- |
| Clinical state | 0.736 |
| Overall dopaminergic burden | 0.736 |
| Magnitude-only imaging | 0.736 |
| Direction-aware imaging | 0.823 |

Table 18: Earlier matched-profile audit: no clear association between secondary-profile distance and future divergence. Intervals are pair-bootstrap 95% intervals; P values are two-sided permutation tests. These related exploratory screens are not independent replications.

| Coverage / distance caliper | People | Pairs | Pearson r (95% CI) | P |
| --- | --- | --- | --- | --- |
| Complete / 75th percentile | 376 | 122 | -0.059 (-0.209, 0.104) | 0.515 |
| Complete / 50th percentile | 376 | 77 | -0.092 (-0.265, 0.115) | 0.432 |
| MoCA and CSF / 75th | 380 | 122 | -0.063 (-0.208, 0.095) | 0.490 |
| SAA / 75th | 616 | 204 | 0.038 (-0.104, 0.186) | 0.591 |

In the complete-profile comparison, adjusting for residual baseline matching distance gave a standardized slope of -0.059 (95% CI -0.211 to 0.105). The null result means that visible secondary biomarker differences cannot presently explain the illustrated patients’ different future side balance. This boundary is compatible with a reproducible directional DaT–motor association: preserving one coordinate is not equivalent to establishing a complete mechanistic account of disease progression.

#### S8. Original regional measurements behind the patient displays

The four preselected cases each link to one original scan row in the landmark month and one selected clinical examination. Regional values below reproduce the saved means and ratios; no new subject selection or model fitting was performed. Putamen is the primary directional coordinate, while caudate provides secondary regional context.

| Case | Region | Left SBR | Right SBR | Mean SBR | Signed ratio |
| --- | --- | --- | --- | --- | --- |
| A | Putamen | 0.62 | 1.07 | 0.845 | 0.2663 |
| A | Caudate | 1.82 | 2.19 | 2.005 | 0.0923 |
| B | Putamen | 1.26 | 0.56 | 0.910 | -0.3846 |
| B | Caudate | 2.43 | 2.19 | 2.310 | -0.0519 |
| C | Putamen | 0.58 | 0.57 | 0.575 | -0.0087 |
| C | Caudate | 1.19 | 1.73 | 1.460 | 0.1849 |
| D | Putamen | 0.15 | 0.76 | 0.455 | 0.6703 |
| D | Caudate | 2.12 | 2.28 | 2.200 | 0.0364 |

The regional putamen–caudate visualization appears earlier in Figure 6; the table above reproduces the same source measurements for the Supplementary Information record.

Case C’s paired motor sums (left 1, right 11) give clinical laterality 0.8333 despite minimal putamen direction. Its caudate direction is concordant, but substituting caudate after viewing a case would change the hypothesis and is not done. In Case D, left/right motor sums 12/9 give -0.1429 despite positive putamen asymmetry. The separate regional and clinical findings remain visible; an attractive anatomical narrative cannot erase disagreement. Cases A/B also differ in total motor burden (31/12), so their contrast is not a controlled comparison of otherwise equivalent disease states.

